# The role of intestine in metabolic dysregulation in murine Wilson disease

**DOI:** 10.1101/2023.01.13.524009

**Authors:** Gaurav V. Sarode, Tagreed A. Mazi, Kari Neier, Noreene M. Shibata, Guillaume Jospin, Nathaniel H.O. Harder, Marie C. Heffern, Ashok K. Sharma, Shyam K. More, Maneesh Dave, Shannon M. Schroeder, Li Wang, Janine M. LaSalle, Svetlana Lutsenko, Valentina Medici

## Abstract

**Background and aims:** Major clinical manifestations of Wilson disease (WD) are related to copper accumulation in the liver and the brain, and little is known about other tissues involvement in metabolic changes in WD. *In vitro* studies suggested that the loss of intestinal ATP7B could contribute to metabolic dysregulation in WD. We tested this hypothesis by evaluating gut microbiota and lipidome in two mouse models of WD and by characterizing a new mouse model with a targeted deletion of *Atp7b* in intestine.

**Methods:** Cecal content 16S sequencing and untargeted hepatic and plasma lipidome analyses in the Jackson Laboratory toxic-milk and the *Atp7b* null global knockout mouse models of WD were profiled and integrated. Intestine-specific *Atp7b* knockout mice (*Atp7b*^ΔIEC^) was generated using B6.Cg-Tg(Vil1-cre)997Gum/J mice and *Atp7b*^Lox/Lox^ mice, and characterized using targeted lipidome analysis following a high-fat diet challenge.

**Results:** Gut microbiota diversity was reduced in animal models of WD. Comparative prediction analysis revealed amino acid, carbohydrate, and lipid metabolism functions to be dysregulated in the WD gut microbial metagenome. Liver and plasma lipidomic profiles showed dysregulated tri- and diglyceride, phospholipid, and sphingolipid metabolism in WD models. When challenged with a high-fat diet, *Atp7b^ΔIEC^* mice exhibited profound alterations to fatty acid desaturation and sphingolipid metabolism pathways as well as altered APOB48 distribution in intestinal epithelial cells.

**Conclusion:** Coordinated changes of gut microbiome and lipidome analyses underlie systemic metabolic manifestations in murine WD. Intestine-specific ATP7B deficiency affected both intestinal and systemic response to a high-fat challenge. WD is a systemic disease in which intestinal-specific ATP7B loss and diet influence phenotypic presentations.

## INTRODUCTION

Wilson disease (WD) is a rare disease characterized by a spectrum of severity of hepatic and neuro-psychiatric clinical manifestations. The traditional and established view of WD pathogenesis is rooted in the role of the copper-transporting ATPase beta (ATP7B) as mutations in the gene are the cause. Because of *ATP7B* disease-causing variants, the function of ATP7B is lost or greatly diminished, and copper accumulates in hepatocytes, leading to cellular and systemic metabolic alterations including oxidative stress, mitochondrial changes, and perturbed lipid and methionine metabolism^1–3^. Previous untargeted metabolomic profiling revealed altered phospholipid levels in patients with WD, corroborated by findings in a mouse model of WD^4, 5^. Fatty liver is a frequent manifestation of WD and it occurs mostly in absence of obesity and other features of metabolic syndrome. Diet has emerged as a key modulator of WD pathogenesis. An obesogenic diet with high fructose corn syrup and high-sucrose content induces alterations in both lipid profiles and parameters of copper metabolism^6^. In a rat model of WD, feeding a high-calorie Western diet exacerbated mitochondria dysfunction, hepatic steatosis, and liver damage^3^.

Murine models of WD play a crucial role in the understanding of its pathogenesis as they reliably accumulate copper and develop hepatic histological manifestations resembling human liver disease. The *Atp7b*^-/-^ mouse model has metabolic phenotypes, including impaired cholesterol synthesis pathways similar to those described in patients with WD ^7^. *Atp7b* null mice maintained on different backgrounds (hybrid or C57Bl/6) consistently showed similar lipid metabolism changes^8^, despite different rates of copper accumulation. The Jackson Laboratory toxic-milk mouse C3He-Atp7b^tx-J^/J (tx-j), characterized by a spontaneous mutation of *Atp7b* and hepatic copper accumulation, also has altered lipid metabolism that partially overlaps with findings in WD patients^9^.

Although the primary sites of WD clinical manifestations are the liver and the brain, growing evidence indicates multiple organs are affected, including the intestine, heart, and kidneys. However, it remains unknown whether the involvement of extra-hepatic organs is independent of liver defects or secondary to hepatic copper accumulation. The ATP7B tissue-specific roles are also largely unknown. A role of ATP7B in intestinal epithelial cells (IECs), gut lipid metabolism and chylomicron maturation was demonstrated in a ground-breaking study using intestine-derived organoids from *Atp7b* null global knockout mice on a hybrid C57BL/6 × 129S6/SvEv background (*Atp7b*^-/-^)^10^. Therefore, it is possible the inactivation of ATP7B in extra-hepatic tissues, such as the intestine, directly contributes to systemic manifestations of WD, independently of hepatic copper accumulation and hepatic pathological changes.

Given that both copper and lipid metabolism occur in the liver and IECs, it is likely that dietary factors and the gut microbiome have an effect on systemic WD energy metabolism with consequent impacts on the risk of developing hepatic steatosis and liver damage.

In addition, initial studies in patients with WD indicate reduced alpha diversity of their gut microbiome^11^ and our untargeted metabolomic profile in WD revealed a complex metabolic interaction involving dietary and microbiome-derived metabolites or metabolites for which plasma levels are affected by microbiome composition^9^.

The goal of the present study is to profile and integrate gut microbiome and lipidomic analyses in two mouse models of WD. Results were mechanistically validated in a newly generated intestine-specific *Atp7b* knockout mouse model (*Atp7b*^ΔIEC^), designed to study the role of intestine ATP7B on lipid and copper metabolism independent of hepatic copper accumulation and liver disease.

## MATERIALS AND METHODS

### *Atp7b^ΔIEC^* model generation

*Atp7b*^ΔIEC^ mice were generated by the University of California Davis Mouse Biology Program using B6.Cg-Tg(Vil1-cre)997Gum/J mice from the Jackson Laboratory and *Atp7b^Lox/Lox^* mice^12^ kindly provided by Dr. Svetlana Lutsenko (Johns Hopkins University). Cre-mediated removal of a 1.6-kb fragment in exon 2 results in IEC-specific *Atp7b* inactivation. Gene and protein expression were performed to confirm IEC-specific *Atp7b* inactivation (Figure S1).

### Experimental animals

All protocols were approved by the UC Davis Institutional Animal Care and Use Committee and follow the National Research Council’s Guide for the Care and Use of Laboratory Animals. The *Atp7b* null global knockout on a C57Bl/6 background was generated as previously described^12^; this model will hereafter be referred to as “KO” to distinguish it from the hybrid background model. The KO model on a C57Bl/6 background for the present study as the development of liver pathology in these mice resembles the natural history of the tx-j mouse. KO, *Atp7b*^ΔIEC^, tx-j, and C3HeB/FeJ (C3H) colonies were bred and maintained on the UC Davis campus in standard plastic shoebox cages with Teklad TEK-Fresh bedding (Envigo, Madison, WI) and nesting enrichment material under the following conditions: 20–23°C, 40%–65% relative humidity, 14 h light/10 h dark light-cycle, and *ad libitum* LabDiet 5001 (PMI, St. Louis, MO) and deionized water. Additional study details and experimental methods can be found in Supplemental Material.

## RESULTS

As shown in Table S3, at 16 weeks of age, tx-j mice presented significantly lower body weight compared to C3H control with normal copper metabolism, as previously described^13^; KO body weights were not different from *Atp7b^+/+^* littermate controls (WT) of the same sex. Female mice of both strains presented lower body weight compared to their male counterparts. Liver per body weight was significantly higher in female tx-j compared with C3H control and male tx-j mice. Mesenteric white adipose tissue per body weight was significantly lower in tx-j mice compared with control but was not different between KO and WT mice.

### Systemic loss of *Atp7b* impacts gut microbiome

To examine changes in the intestinal microbiome associated with *Atp7b* mutation independent of advanced liver disease, a microbiome profile was conducted on 16-week-old mice (see Figure S2 for study design) because at this age both tx-j and KO mice already accumulate copper in the liver^8, 14–16^ but do not yet show significant hepatic steatosis or fibrosis (Figure S3).

Cecal content 16S sequencing revealed decreased alpha diversity of the tx-j microbiome when compared with C3H control, as estimated by observed species index (p=0.002), Chao1 index (p=0.003), and Shannon index (p=0.001) (Figure 1A). Beta diversity analysis (Figure 1B) demonstrated significant partitioning of bacterial communities (F=4.9355; R-squared=0.1499; p<0.002).

**Figure 1.**
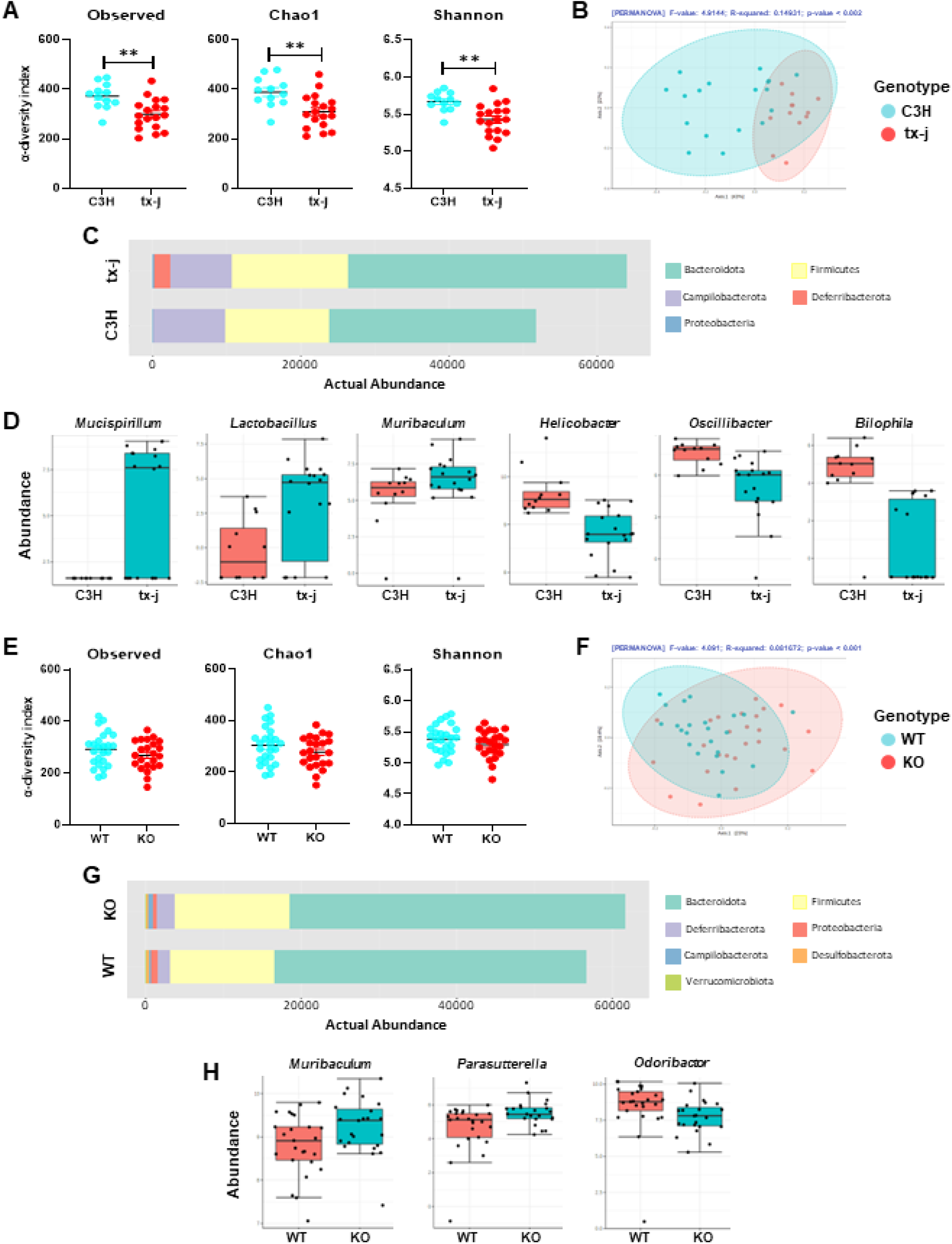
Composition and structure of cecal content microbiota profiles in 16-week-old tx-j vs C3H and KO vs WT mice. A, E: Alpha-diversity analysis shown as Observed species richness, Chao1 index, and Shannon index. Scatter plots are shown as the mean ± SD (**p < 0.01). B, F: Microbial beta-diversity was accessed by NMDS analysis based on Bray-Curtis dissimilarity index by permutational multivariate analysis of variance. C, G: Microbial abundance at the phylum level. D, H: Box plots representing the relative abundance of the top significantly altered bacteria at the genus level.

When examining differences in gut microbiota composition at the phylum and genus levels in tx-j vs C3H, a total of five bacterial phyla (Bacteroidota, Campilobacterota, Deferribacterota, Firmicutes, and Proteobacteria) were dominant (Figure 1C).

In the taxonomic profile for tx-j vs C3H, the operational taxonomic units were assigned to prevalent microbiome components of the phyla Bacteroidota (53% vs 46%), Firmicutes (32% vs 39%), Campilobacterota (10% vs 14%), and Deferribacterota (4% vs 1%) (Figure S4). A clustered heat map showing the variation of taxonomic abundance at the genus level by genotype and sex in tx-j vs C3H is presented in Figure S6.

The proportions of tx-j gut bacteria at the genus level were very distinct from those in C3H control mice. A total of 26 genera were identified of which 10 showed significant abundance differences according to classical univariate analysis (p<0.05) (Table S4). The relative abundances of *Mucispirillum, Lactobacillus, Muribaculum, Rikenellaceae_RC9_gut_group*, and *Odoribacter* were significantly higher in tx-j mice compared with C3H. *Helicobacter, Oscillibacter, Bilophila, Blautia*, and *Alistipes* were less abundant, however, did not retain significance with FDR correction. The top 6 differential genera are shown in Figure 1D.

### Mice genetic background modulates microbiome response

On 16S sequencing, in terms of alpha and beta diversity, KO showed similar trends compared to tx-j mice control but differences were not statistically significant (Figure 1E and 1F). In KO vs WT groups, seven bacterial phyla (Bacteroidota, Campilobacterota, Deferribacterota, Firmicutes, Proteobacteria and Verrucomicrobiota) were dominant (Figure 1G). Twenty-three genera were identified with only *Muribaculum*, *Parasutterella*, *Bacteroides*, and *Desulfovibrio* presenting significantly higher abundances in KO mice compared with WT, and *Odoribacter* less abundant (Figure 1H). Bacterial phyla analysis did not show differences between KO and WT mice (Figure S5).

### Functional metagenome predicts involvement of energy metabolism

To understand the microbiome’s functional composition, we performed a comparative prediction analysis of the functional metagenome using PICRUSt. MicrobiomeAnalyst was used to map the abundance profile for functional analysis of microbiota. Unsupervised clustering by principal component analysis (PCA) showed the tx-j microbiota profile separated from that of C3H mice and showed 44.8% of variance between the groups (Figure S7). Moreover, classical univariate analysis identified 1,957 significant KEGG orthology groups between tx-j and C3H mice. According to KEGG metabolic network pathway analysis and as indicated in Table S6, the observed KEGG orthology shows primary involvement of energy metabolism including amino acid biosynthesis, metabolism of branched-chain amino acids, pyruvate metabolism and the tricarboxylic acid cycle. Compatibly with the less pronounced differences in cecal microbiome patterns in KO vs WT, less marked associations between microbiome and energy metabolism were found in the KO mice; regardless, associations were in the same direction as tx-j mice confirming the role of genotype on influencing both microbiome and energy metabolism pathways (Table S6).

### Lipidome analyses of liver and plasma identify specific lipid changes in mouse models of Wilson disease

To examine the effect of ATP7B deficiency on lipid metabolism independently from liver disease, we compared liver and plasma lipidomic profiles in 16-week-old tx-j and KO mouse models when fed a standard chow diet. PCA indicated distinct clustering and separation between tx-j and C3H liver (Figure 2A) and plasma (Figure 2B). When examining enrichment analysis in both liver (Figure 2C) and plasma (Figure 2D), lipid classes in tx-j mice showed lower triglycerides (TGs), diglycerides (DGs), phospholipids (PLs), and sphingomyelins (SMs) along with higher ceramides compared with C3H control mice. Plasma levels of unsaturated and saturated fatty acids were lower in tx-j mice compared with C3H control.

**Figure 2.**
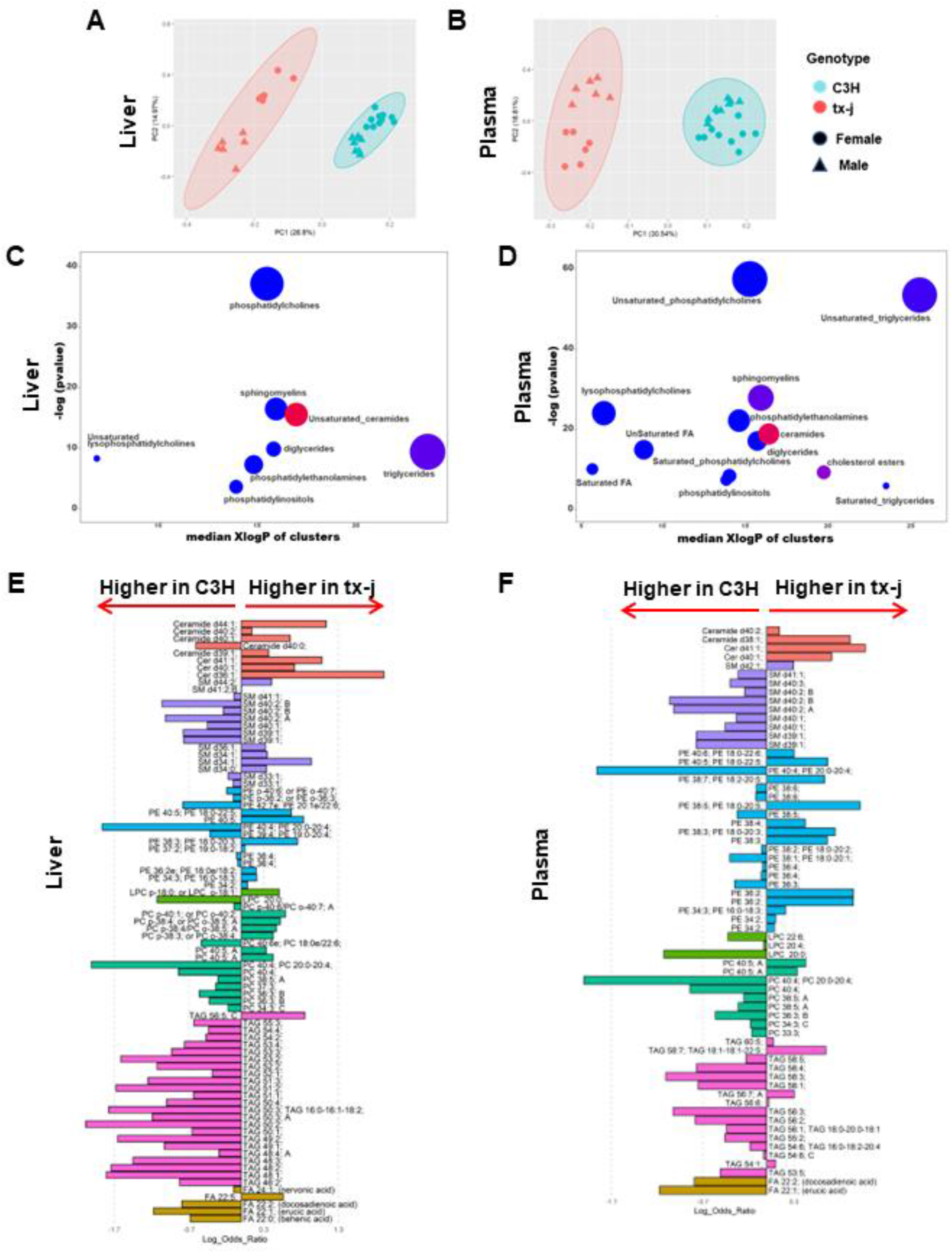
Lipidomic profiles in liver and plasma of tx-j mice compared with C3H control. A, B: Principal component analysis based on the lipid profiling of tx-j mice and C3H control in liver and plasma. C, D: Chemical similarity enrichment analysis (ChemRICH) and enrichment statistics plot for tx-j vs C3H mice in liver and plasma. Each cluster represents an altered chemical class of metabolites (p<0.05). Cluster size represents the total number of metabolites. Cluster color represents the directionality of metabolite differences: red, higher in tx-j mice; blue, lower in tx-j mice. Colors in between refer to a mixed population of metabolites manifesting both higher and lower levels in tx-j mice when compared with C3H. The x-axis represents the cluster order on the chemical similarity tree. The y-axis represents chemical enrichment p-values calculated using a Kolmogorov–Smirnov test. E, F: Linear discriminant analysis effect size (LEfSe) plot representing the logarithm of the ratios of the average levels of statistically different lipids between tx-j and C3H mice in liver and plasma. A positive log ratio bar indicates higher lipid levels in tx-j mice; a negative log ratio bar indicates higher levels in C3H mice. Colors represent lipids in the same class.

The hepatic lipidome in tx-j mice was particularly characterized by changes in unsaturated lipid classes, including lower levels of several triacylglycerides (TAGs) and phosphatidylcholines (PCs) with an altered phosphatidylethanolamine (PE) profile, indicating dysregulated PL metabolism. The sphingolipid profile was affected by higher ceramides and altered SMs. In addition, hepatic very-long-chain fatty acid (VLCFA) pathways were aberrant (Figure 2E). Likewise, in plasma, we observed altered TAGs and lower levels of many PCs and lysophosphatidylcholines (LPCs), with PE profile changes. Higher ceramides, lower SMs, and altered VLCFAs were also observed (Figure 2F). A list of all significantly altered metabolites is presented in Supplemental Material 2 (Excel file).

Consistent with findings from the tx-j enrichment analysis and compatibly with less pronounced microbiome pattern differences, 16-week-old KO mice showed less separation in lipidomic profiles (Figure 3A and 3B) but similar trends in liver and plasma lipidomic patterns compared to WT mice (Figure 3C, D, E, and F).

**Figure 3.**
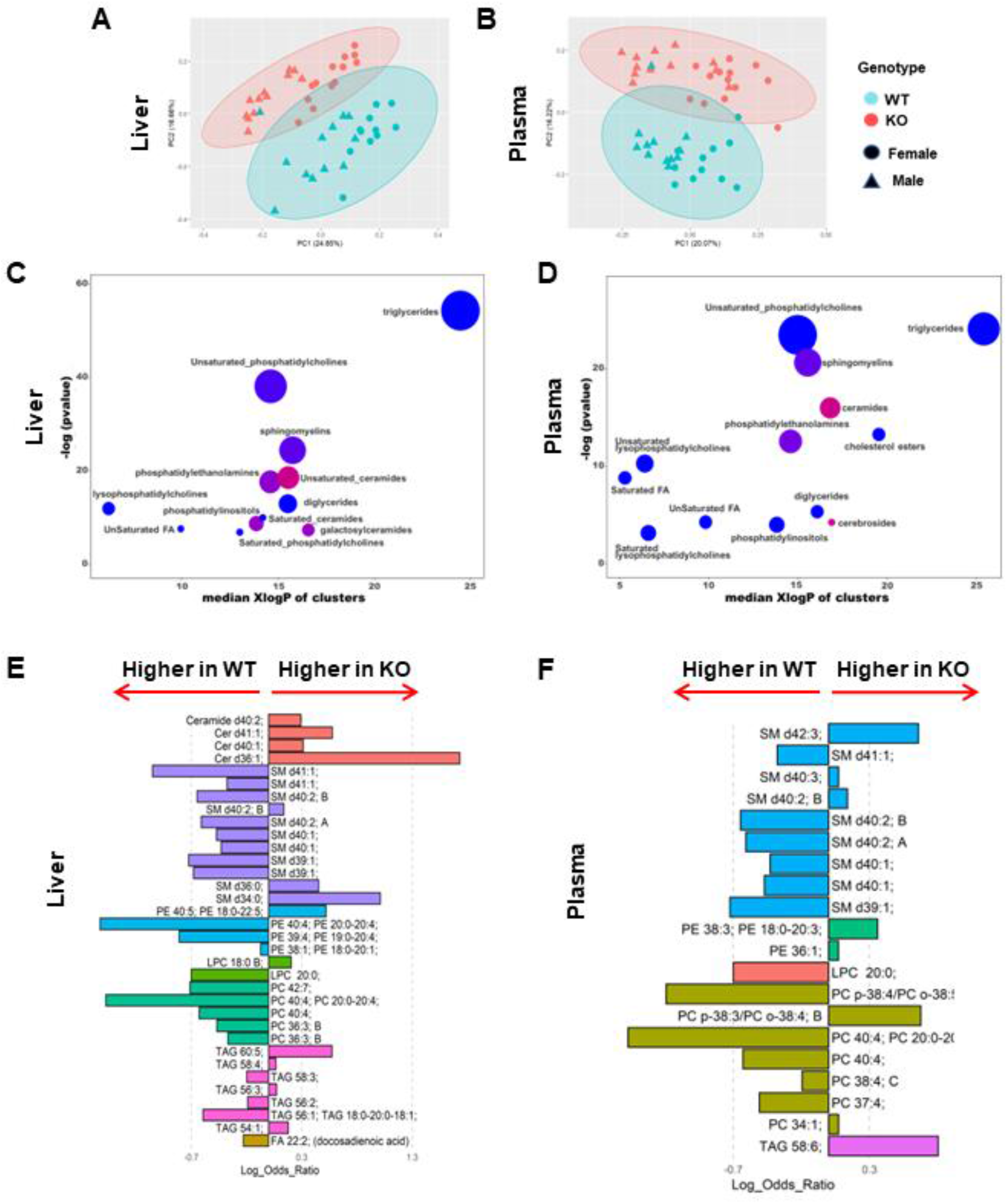
Lipidomic profiles in liver and plasma of KO mice compared with WT control. A, B: Principal component analysis based on the lipid profiling of KO mice and WT control in liver and plasma. C, D: Chemical similarity enrichment analysis (ChemRICH) and enrichment statistics plot for KO vs WT mice. Each cluster represents an altered chemical class of metabolites (p<0.05). Cluster size represents the total number of metabolites. Cluster color represents the directionality of metabolite differences: red, higher in KO mice; blue, lower in KO mice. Colors in between refer to a mixed population of metabolites manifesting both higher and lower levels in KO mice when compared with WT. The x-axis represents the cluster order on the chemical similarity tree. The y-axis represents chemical enrichment p-values calculated using a Kolmogorov–Smirnov test. E, F: Linear discriminant analysis effect size (LEfSe) plot representing the logarithm of the ratios of the average levels of statistically different lipids between KO and WT mice in liver and plasma. A positive log ratio bar indicates higher lipid levels in KO mice; a negative log ratio bar indicates higher levels in WT mice. Colors represent lipids in the same class.

### Changes in gut microbiome correlate with changes in hepatic lipidome illustrating contribution to Wilson disease metabolic dysregulation

We used the tx-j model to determine the association of microbiome profiles with hepatic lipidome. Specifically, we focused on microbiota genera exhibiting significant abundance and lipid classes that showed significant differences in tx-j compared with C3H control (Figure 4A and B). With these criteria, we included eight genera (*Mucispirillum, Lactobacillus, Muribaculum, Odoribacter, Bilophila, Oscillibacter, Helicobacter, Blautia*) and selected hepatic TAGs, PLs, ceramides, SMs, and VLCFAs for correlation analysis (Supplemental Material 3 Excel file). Similar criteria were used with lipid-microbe correlation analysis in KO vs WT (Figure 4C and 4D) and included 5 genera (*Muribaculum*, *Odoribacter*, *Desulfovibrio*, *Parasutterella*, *Bacteroides*).

**Figure 4.**
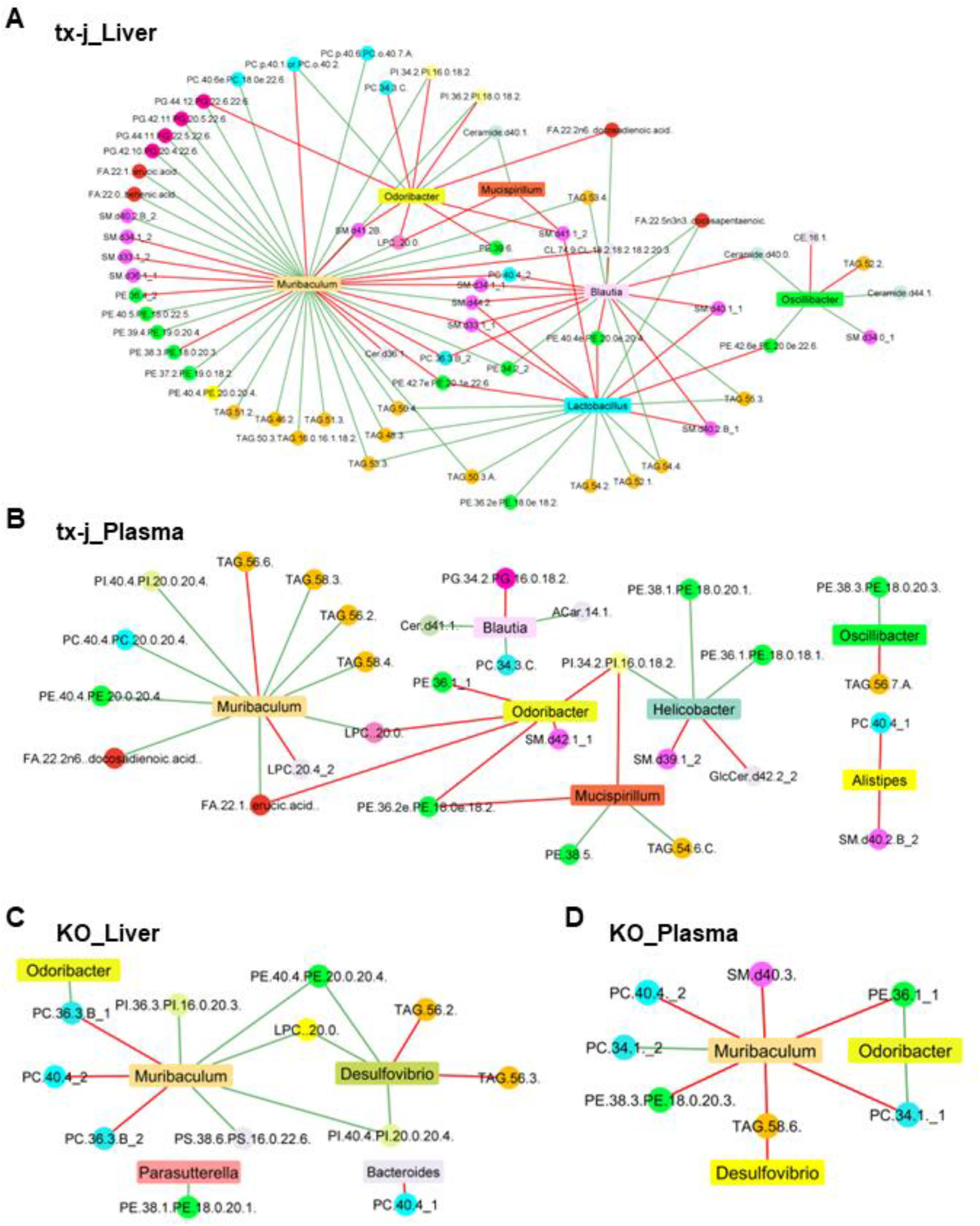
Microbe-metabolite network plots for liver and plasma of tx-j and KO mice. A, B: Microbe-metabolite relational networks showing all significant bacterial genus-lipid metabolite pairs (p≤0.05) between liver and plasma of tx-j and C3H mice. C, D: Microbe-metabolite relational networks showing all significant bacterial genus-lipid metabolite pairs (p≤0.05) between liver and plasma of KO and WT mice. Compositionally corrected correlations (calculated using CCREPE function in R) between microbial taxa and metabolite intensities were used to generate the network plots in CytoScape. In the network plot, the color of the lines represents positive (green) or negative (red) correlations. Nodes of the network represent microbial taxa (rectangle) and lipids (circle); lipid nodes with the same color belong to the same lipid class.

Many TAG species, such as TAG(46-58:1-5), were positively correlated with *Mucispirillum, Lactobacillus*, and *Blautia* (r=0.4-0.6, FDR-adjusted p<0.05). *Muribaculum, Lactobacillus, Odoribacter*, and *Blautia* were negatively associated with many PLs including PC(36-40:3-4) and PE(36-42:3-7); however, *Mucispirillum* was positively associated with many PLs including PC(40:4-6) and PE(34-40:2-5) (r=0.4-0.7, FDR-adjusted p<0.05). *Mucispirillum, Lactobacillus*, and *Blautia* were found to be negatively correlated with several SMs, including SM(33-41:1-2) (r=0.4-0.7, FDR-adjusted p<0.05). *Mucispirillum, Odoribacter*, and *Oscillibacter* were positively correlated with some ceramides, including Cer(d40-44:0-1) (r=0.4-0.7, FDR-adjusted p<0.05). Together, the correlations observed between *Mucispirillum, Muribaculum Lactobacillus*, and lipids classes, such as PLs and SMs, are in accordance with the direction of change observed in lipidomic profiles of tx-j and KO mice. Using a systems-based approach to explore the liver data in more depth, we performed a weighted gene correlation network analysis of the microbiome and liver lipidome data of tx-j and KO mice. Through this analysis, a network of liver lipids for both tx-j and KO mice was constructed and organized in color-coded modules (Figure S8 and related table with lipid membership in each module). With this unbiased approach, we similarly identified major lipid modules associated with WD in both tx-j and C3H and KO vs WT mouse genotypes. Next, we studied genera-module associations, focusing on *Mucispirillum* given its previously described central association with metabolic changes characteristic of fatty liver. Liver lipid modules midnight blue, green yellow, green, and black were all significantly associated with *Mucispirillum*, particularly pointing to PC and LPC as identified associated lipids (Figure S9).

### Lipid metabolism is altered in intestinal epithelial cells of mice with targeted *Atp7b* deletion

After the a 8-day high-fat diet (HFD) challenge, the intestinal epithelium in newly generated *Atp7b^ΔIEC^* mice presented an APOB48 cytosolic distribution resembling the aberrant distribution previously described in organoids derived from *Atp7b^-/-^* mouse IECs^10^ (Figure 5Ad). IECs isolated from KO and *Atp7b*^ΔIEC^ mice also presented similar patterns regarding increased levels of factors central in lipid metabolism, particularly lipid metabolism associated with copper accumulation, including phosphorylated-AMP-activated protein kinase and peroxisome proliferator-activated receptor alpha (PPARα) and gamma (PPARγ) (Figure 5B and 5C).

**Figure 5.**
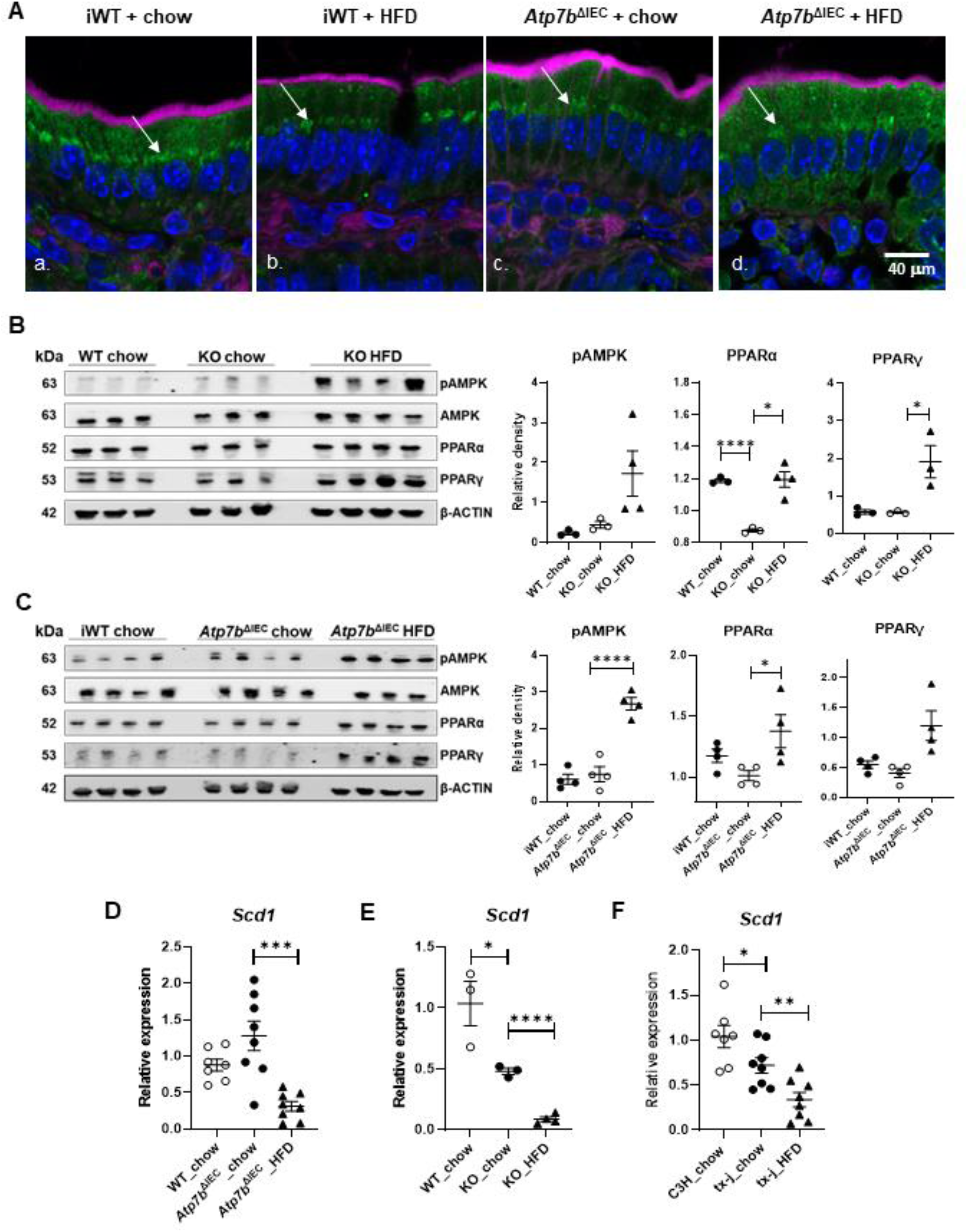
Lipid metabolism in intestinal epithelial cells of mice with abnormal copper metabolism, fed chow or HFD. A: Immunofluorescence for APOB48 in the intestine of iWT and *Atp7b*^ΔIEC^ mice fed chow or HFD. APOB48 (green) presented similar peri-nuclear distribution (arrows) in iWT mice, regardless of the dietary fat content (a, b), and in *Atp7b*^ΔIEC^ mice fed chow (c). However, after HFD challenge in *Atp7b*^ΔIEC^ mice, APOB48 presented a diffuse cytosolic distribution indicating an impairment of chylomicron processing within the IECs (d). B, C: Immunoblots and densitometry analyses of metabolic regulators (AMPK, PPARα, and PPARγ) in 9-week-old IECs from KO and *Atp7b^ΔIEC^* mice and their respective controls fed chow or HFD. Immunoblot densitometry analyses are normalized to β-ACTIN; data are represented as means ± SEM with statistical significance determined by Student’s t-test (*p < 0.05, and ****p < 0.0001). D, E, F: qPCR transcript levels of stearoyl-CoA desaturase 1 (*Scd1*) measured in livers of 9-week-old *Atp7b*^ΔIEC^, KO, and tx-j mice with their respective controls (iWT, WT, and C3H) fed chow or 60% kcal fat diet (HFD). Transcript levels are normalized to *Rplp0;* data are represented as means ± SEM with statistical significance determined by Student’s t-test (*p < 0.05, **p < 0.01, ***p < 0.001, and ****p < 0.0001).

### Targeted lipidome analysis of liver *Atp7b*^ΔIEC^ demonstrates contribution of intestinal ATP7B to systemic metabolic dysregulation in Wilson disease

*Atp7b*^ΔIEC^ mice body weights and liver per body weight ratios were not different compared to Lox/Lox:Cre-littermates as controls (iWT) at 9 weeks of age (Table S7). However, liver histology of chow-fed *Atp7b^ΔIEC^* mice presented hepatocyte cytosolic glycogenosis and vacuolization (Figure S10). To understand the effect of intestine ATP7B removal on lipid metabolism independent from copper accumulation in the liver, we examined liver and plasma-targeted lipidomic profiles in *Atp7b^ΔIEC^* mice on chow compared with their Lox/Lox:Cre-littermates as controls (iWT). We observed subtle changes highlighting increased liver TG, DGs, and unsaturated FAs. In plasma, *Atp7b^ΔIEC^* mice showed higher levels of lysoPCs and lysoPEs and reduced SM and ceramides (Figure S11).

We examined the systemic effects of the 8-day course of HFD on *Atp7b^ΔIEC^* liver and plasma targeted lipidomic profiles at 9 weeks of age. Compatibly with the short course of HFD, body weights and liver per body weight ratios (Table S7) showed no differences within genotypes after short-course HFD. Liver histology of HFD-fed *Atp7b^ΔIEC^* mice presented cytosolic glycogenosis and vacuolization similarly to chow-fed mice of the same genotype (Figure S10). For the targeted lipidomic analysis, we elected to analyze differentially abundant lipid classes from the untargeted analysis comparing 16-week-old KO and WT mice.

*Atp7b*^ΔIEC^ mice challenged with HFD showed distinct clustering and separation from WT and chow-fed *Atp7b*^ΔIEC^ liver and plasma as indicated by clustering (Figure 6A) and PCA analysis (Figure 6B-C). Specifically, the hepatic lipidome profile presented a dramatic increase in TAGs and DGs, and lower acylcarnitines, HFD also resulted in altered ceramides and increased SMs as well as a dysregulated PL profile with lower levels of many PCs and PEs, and altered lysoPCs and lysoPEs (Figure 6D). In plasma, HFD resulted in lower TAGs and higher acylcarnitines, ceramides, and SMs (Figure 6E). Of note, hepatic gene transcript analysis of *Scd1*, a central player in the fatty acid desaturation pathway, were not different in *Atp7b*^ΔIEC^ mice on chow diet compared with iWT and they were significantly reduced after HFD (Figure 5A).

**Figure 6:**
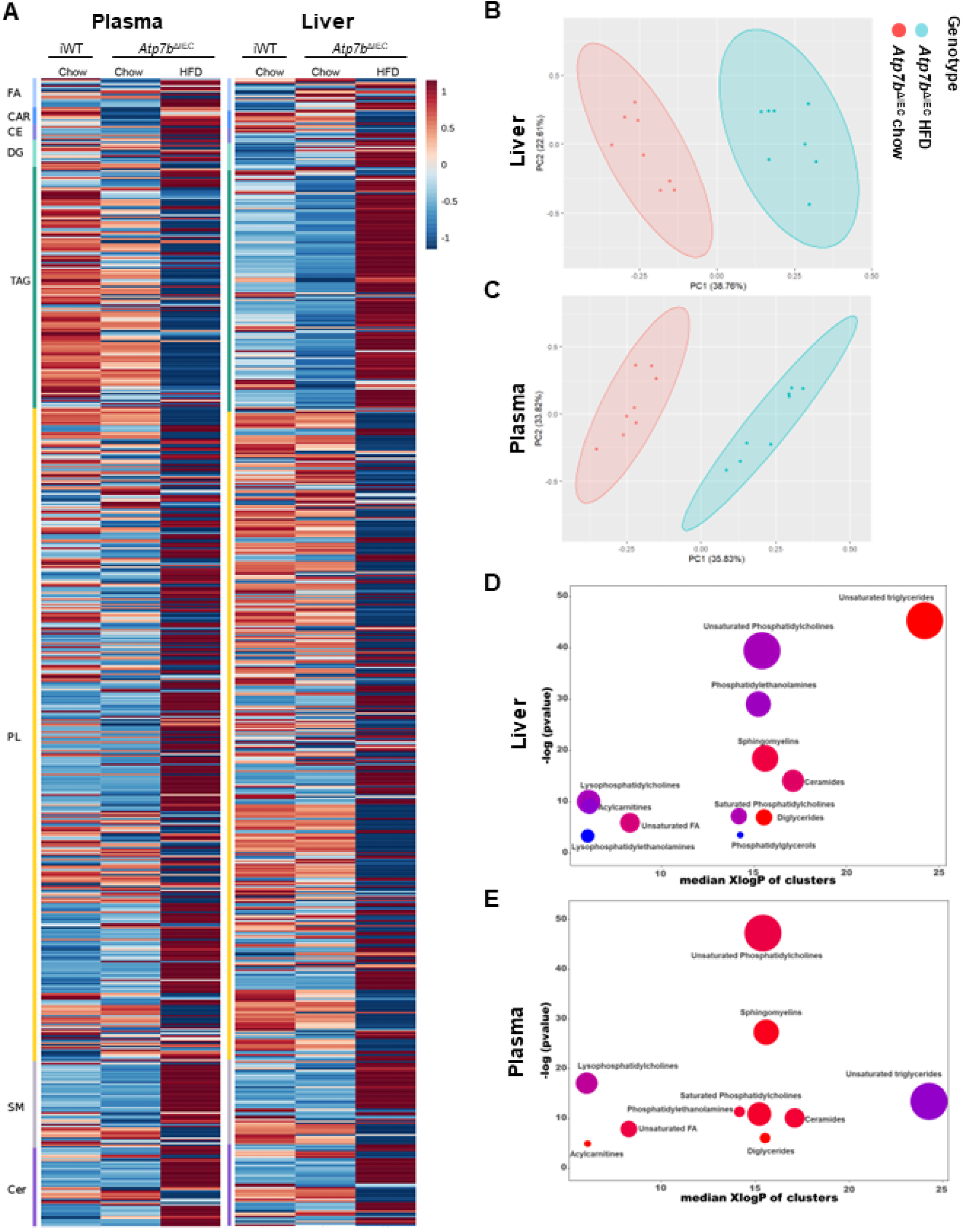
Lipidomic profiles in liver and plasma of intestine-specific *Atp7b* knockout mice (*Atp7b*^ΔIEC^) fed chow or HFD. A: Heat map depicting plasma and liver lipidomic profiles in *Atp7b*^ΔIEC^ mice compared to their Lox^+/+^/Cre^-^ littermate control (iWT). Group means are shown using normalized data with direction and degree of change indicated by color and intensity, respectively (red = increase, blue = decrease). CAR, acylcarnitines; CE, cholesterol ester; Cer, ceramide; DG, diglyceride; FA, fatty acid; HFD, high fat diet; PL, phospholipid; SM, sphingomyelin; TAG, triacylglyceride. B, C: Principal component analysis based on the lipid profiling of *Atp7b*^ΔIEC^ mice fed chow or HFD in liver and plasma. D, E: Chemical similarity enrichment analysis (ChemRICH) and enrichment statistics plot for *Atp7b*^ΔIEC^ HFD vs *Atp7b*^ΔIEC^ chow mice in liver and plasma. Each cluster represents an altered chemical class of metabolites (p < 0.05). Cluster size represents the total number of metabolites. Cluster color represents the directionality of metabolite differences: red, higher in *Atp7b*^ΔIEC^ HFD mice; blue, lower in *Atp7b*^ΔIEC^ chow mice. Colors in between refer to a mixed population of metabolites manifesting both higher and lower levels in *Atp7b*^ΔIEC^ HFD when compared with *Atp7b*^ΔIEC^ chow. The x-axis represents the cluster order on the chemical similarity tree. The y-axis represents chemical enrichment p-values calculated using Kolmogorov–Smirnov test.

We then examined the effects of HFD on liver and plasma targeted lipidomic profiles in iWT mice with normal copper metabolism and compared to WT mice on the same diet (Figure 7A). Mice were characterized by reduced saturated and unsaturated TGs and ceramides as well as extensive alterations in SM and PE in the liver (Figure 7C and Figure 7E). Plasma unsaturated TGs were also markedly reduced with lower ceramide levels (Figure 7D and Figure 7F). When studying the effects of HFD on lipidomics in KO mice, PCA indicated marginal separation between chow and HFD KO mice in both liver (Figure S12A) and plasma (Figure S12B). Compared to chow, the KO hepatic lipidome showed increased DGs and a trend in increased TAGs (not FDR significant) on HFD (Figure S12C) in a similar direction compared to *Atp7b^ΔIEC^* mice. Whereas no changes in hepatic ceramides were observed, the PL profile was found altered with lower levels of many PCs (Figure S12E). In plasma, the HFD challenge resulted in a decrease in several TAGs along with altered PL profiles and increased levels of ceramides and SMs (Figure S12F). Similar to *Atp7b*^ΔIEC^ mice, *Scd1* transcript levels (Figure 5E and Figure 5F), were further downregulated after the HFD challenge in both KO and tx-j mice.

**Figure 7:**
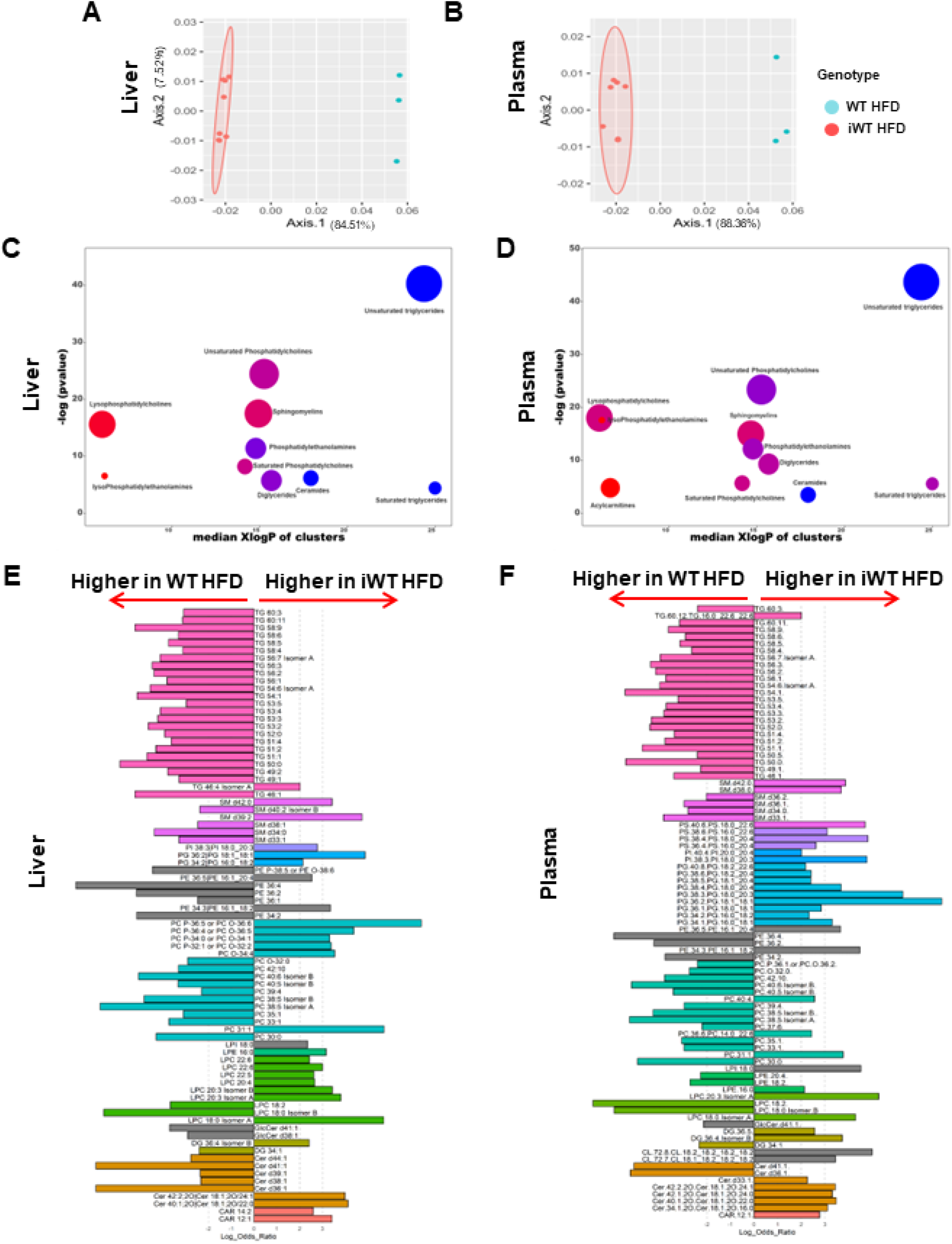
A, B: Principal component analysis based on the lipid profiling of WT and iWT mice fed HFD in liver and plasma. C, D: Chemical similarity enrichment analysis (ChemRICH) and enrichment statistics plot for iWT HFD vs WT HFD mice. Each cluster represents an altered chemical class of metabolites (p < 0.05). Cluster size represents the total number of metabolites. Cluster color represents the directionality of metabolite differences: red, higher in iWT HFD mice; blue, lower in iWT HFD mice. Colors in between refer to a mixed population of metabolites manifesting both higher and lower levels in iWT HFD when compared with WT HFD. The x-axis represents the cluster order on the chemical similarity tree. The y-axis represents chemical enrichment p-values calculated using a Kolmogorov-Smirnov test. E, F: Linear discriminant analysis effect size (LEfSe) plots representing the logarithm of the ratios of the average levels of statistically different lipids between iWT HFD and WT HFD mice in liver and plasma. A positive log ratio bar indicates higher lipid levels in iWT HFD; a negative log ratio bar indicates higher levels in iWT HFD mice. Colors represent lipids in the same class.

## DISCUSSION

We conducted an integrated analysis of the microbiome and lipidomic profile in animal models of WD and further investigated the combined effects of intestine-specific *Atp7b* deficiency and challenge with HFD, resulting in several novel findings. First, global *Atp7b* deficiency was characterized by lower microbiome diversity, as shown in WD mouse models compared to control mice, with patterns highlighting phyla and genera previously associated with liver disease and fatty liver in the general population. Second, global *Atp7b* deficiency was associated with altered lipidomic profiles in two mouse models of WD, specifically dysregulated hepatic fatty acid desaturation and SLs that are independent of severe liver disease. Third, alterations in the microbiome and lipidomic profiles of WD mouse models were highly correlated with microbiome functional analysis supporting the involvement of gut flora composition in WD metabolic complications. Last, we demonstrated a novel mechanism with a potential role in WD pathogenesis, in which intestine-specific *ATP7B* deficiency resulted in a diet-dependent lipidomic dysregulation independent of hepatic copper accumulation and liver disease. Previous studies have shown decreased gut microbiome diversity in WD patients compared to healthy subjects with a higher abundance of Bacteroidetes and lower abundance of Firmicutes, Proteobacteria, and Fusobacteria^17^. Notably, our study supports previous findings in patients and confirmed reduced intestinal flora diversity in animal models of WD, independent of advanced liver disease which influences microbiome composition. In particular, we show genus level changes including a higher abundance of Mucispirillum, Lactobacillus, Rikenellaceae RC9 gut group, Muribaculum, Odoribacter, and Oscillibacter and lower abundance of Helicobacter, Blautia, and Bilophila in tx-j as compared with C3H control mice. Decreased Bifidobacterium and Bacteroides and increased Desulfovibrionaceae, Anaerotruncus, Desulfovibrio, and Mucispirillum were previously reported in high-cholesterol diet-induced NAFLD-hepatocellular carcinoma animals along with decreased 3-indolepropionic acid^18^. We have previously described dysregulated hepatic cholesterol homeostasis^1^ and reduced 3-indolepropionic acid^4^ in WD. The positive correlation between Mucispirillum, Muribaculum, and Odoribacter and hepatic steatosis, oxidative stress, and inflammation in alcohol-associated liver disease^19^, and the association of Bilophila, Paraprevotella, and Suturella with higher hepatic fat content^20^ suggest the implication of these microbiotal species in hepatic pathological manifestations in WD. Findings from functional metagenome analysis suggest that the microbiotal shifts associated with WD are implicated in pathways relevant to energy metabolism including amino acid, carbohydrate metabolism, and lipid metabolism. This result is in agreement with a previous study on dietary copper supplementation in pigs, which showed a decreased abundance of microbiota related to energy metabolism, as lysine and other amino acid biosynthesis, and an increased abundance of microbiota related to amino acid and lipid biosynthesis and metabolism^21^. These data point to correlations between ATP7B defects, microbiome diversity, and WD-associated metabolic misregulations.

Whereas derangements in lipid metabolism are reported in patients with WD^22^, our findings highlight lipidomic dysregulation that is independent of advanced liver disease. These include derangements in fatty acid desaturation and elongation pathways, with lower levels of many very-long-chain fatty acids including behenic acid (22:0), erucic acid (22:1n9), and nervonic acid (24:1n9), which can originate from the denaturation and elongation of palmitic acid (16:0) and stearic acid (18:0). We also observed lower levels of several TAG and PLs with unsaturated double bonds, suggesting a de-enrichment of mono- and polyunsaturated fatty acids in these lipid classes, supported by gene transcript analysis with downregulation of *Scd1*, encoding for delta-9 desaturase. Reduced SCD1 results in decreased endogenous monounsaturated fatty acid production and decreased TG synthesis for lipid storage^23^, which may explain the lean phenotype observed in WD mouse models^5^. While the loss/inhibition of SCD1 was shown to induce favorable metabolic effects, there are mixed findings as the inhibition of hepatic SCD1 was also shown to promote inflammation and stress response^24–26^. Our finding corroborates previous reports from our groups indicating lower activity/transcript levels or impaired function of nuclear receptors like farnesoid X receptor (FXR), liver X receptors (LXRs), heterodimer with retinoid X receptor (RXRα), transcription factors as sterol regulatory element-binding protein (SREBP1c), and peroxisome-proliferator-activated receptor alpha (PPARα) which regulate lipogenic genes including *Scd1* in both animal models and patients with WD^27–32^.

Our approach highlights lipidomic alterations that have been at least in part previously described in WD. We have reported altered SLs metabolism in subjects with WD^9^ and ceramide and sphingomyelin were found to increase in patients with WD^33^. In the current analysis, liver and plasma SLs profiles were altered with higher ceramide and altered sphingomyelin profiles. The involvement of aberrant ceramide metabolism in WD was shown *in vitro* and *in vivo* with increased ceramide levels in WD^34^. The inhibition of ceramide production in a rat model of WD prevented the development of cirrhosis^34^. Evidence from animal studies indicates the role of ceramides in hepatic steatosis, inflammation, hepatocellular apoptosis, and fibrosis^35^. The hepatic and plasma PLs profiles were also found altered, a feature reportedly associated with an increased risk of hepatic steatosis^36^. This finding is consistent with our previous work showing dysregulated serum PL profile in WD patients compared to healthy controls and downregulation of genes involved in PC synthesis and metabolism including phosphatidylethanolamine N-methyltransferase (PEMT) and choline phosphotransferase1^9^.

The acute challenge with a HFD in *Atp7b*^-/-^ further accentuated the downregulation of *Scd1*. This finding aligns with previous evidence indicating the emerging role of diet in WD pathogenesis^3, 6^, and further suggests dysregulations in pathways of fatty acid desaturation as a potential driver for the progression to severe WD presentation. Noteworthy, correlation analysis highlighted the differentially abundant microbiotal species, Mucispirillum, and Lactobacillus were associated with alterations in PL and correlated negatively with many TAG and SMs. Mucispirillum also correlated positively with ceramides. As current evidence indicates that intestinal microbiota modulates lipid metabolism, and demonstrated an association of gut microbiota composition with lipid profiles^37^, the association reported in our study suggests implications of these genera in the observed lipidomic alterations in WD mice.

Importantly, our data demonstrate the role of intestine ATP7B in modulating lipid metabolism that is independent of hepatic copper accumulation or hepatic pathological changes. We show that the intestinal ATP7B transporter deletion resulted in the mislocalization of APOB48 in the intestine and presented a diffuse pattern of cytosolic distribution after the HFD challenge only in *Atp7b*^ΔIEC^ mice. This pattern of distribution likely represents deficits in chylomicron processing and secretion with consequent accumulation of VLDL within the IECs^10^. Ultimately, ATP7B intestine-specific deficiency was associated with diet-specific alterations in plasma and liver lipidomic profiles. Specifically, the HFD challenge in *Atp7b^ΔIEC^* resulted in similar changes to that seen in tx-j and *Atp7b*^-/-^ mice, with downregulation in *Scd*, along with the lower nervonic acid (24:1n9) and palmitoleic acid (16:1n7), and the increase in several PUFAs indicates altered fatty acid desaturation pathways.

HFD in *Atp7b*^ΔIEC^ mice also resulted in decreased plasma TAGs possibly explained by impaired intestinal lipid absorption; increased hepatic TAG and DG possibly due to increased hepatic *de novo* synthesis; along with altered hepatic PLs and SLs. Our findings suggest intestinal ATP7B as a diet-dependent modulator of lipid metabolism and highlight a novel potential mechanism to explain WD pathogenesis and associated dysregulations in lipid metabolism that points to the involvement of extrahepatic organs. The implications of altered intestinal ATP7B and lipidomic profile build upon a pattern of lipid changes after a high-fat diet differs from the pattern previously observed in primary hepatocytes isolated from C57BL/6 mice after a 6-week course of 45% fat diet which was characterized mostly by PC, LPC, and DG increase with TG stability as identified by lipidomic analysis^38^ A sixty percent fat diet provided for 12 weeks to C57BL/6 mice and studied with lipidomic analysis was associated with TG increase and reduction of PC, LPC, and PE^39^ with less marked changes in SM and ceramides.

The *Atp7b*^ΔIEC^ studied here is a new mouse model with still limited metabolic characterization and that was studied only at early time points. Despite this limitation, using a multi-omics approach, we show that ATP7B deficit in two mouse models of WD results in alterations in the microbiome and lipidomic profiles, with dysregulation affecting fatty acid desaturation pathways and SLs metabolism, which were further augmented by HFD. Notably, we observed consistent increases in ceramide levels in the models of WD that are exacerbated by the HFD challenge, pointing to an increased risk of chronic liver damage. These findings support the role of diet and the implication of these pathways in WD progression.

Taken together, our data points to hitherto unstudied interconnectivity of organs and microbiome resulting in changes typically associated with WD pathogenesis and related metabolic changes. Through observations in three mouse models, we demonstrate changes in metabolic pathways associated with changes in lipid profiles and microbiome alterations. The changes highlight the importance of copper homeostasis in systemic energy production.

Our study also highlights a role for extra-hepatic ATP7B in modulating lipid metabolism in WD pathogenesis that was shown to be dependent on diet but independent from liver disease and hepatic copper accumulation. Ultimately, the interplay between copper accumulation and lipid metabolism as shown by -omic profiles underscores the complexity of WD and indicates WD should be considered a systemic disease where energy and lipid metabolism interact at an organ-specific level to modulate the phenotypic presentation.

## Abbreviations

*Atp7b’^-^’*: (*Atp7b* null global knockout mouse on a hybrid background),
*Atp7b^ΔIEC^*: (intestine-specific *Atp7b* knockout mouse),
C3H: (The Jackson Laboratory C3HeB/FeJ),
DGs: (diglycerides),
FXR: (farnesoid X receptor),
LXR: (liver X receptors),
HFD: (high fat diet),
IEC: (intestinal epithelial cell),
iWT: (Lox^+/+^/Cre-; littermate control for *Atp7b^ΔIEC^* mice),
KO: (*Atp7b* null global knockout mouse on a C57Bl/6 background),
LPCs: (lysophosphatidylcholines),
PCs: (phosphatidylcholines),
PCA: (principal component analysis),
PE: (phosphatidylethanolamine),
(PPAR): peroxisome-proliferator-activated receptor,
PLs: (phospholipids),
SCD1: (stearoyl-coenzyme A desaturase 1),
SMs: (sphingomyelins),
SREBP1c: (sterol regulatory element-binding protein),
TAGs: (triacylglycerides),
TGs: (triglycerides),
tx-j: (The Jackson Laboratory toxic-milk mouse),
VLCFA: (very-long-chain fatty acid),
WD: (Wilson disease),
WT: (*Atp7b^+/+^* littermate control for KO mice).

## Author contributions

**Gaurav V. Sarode** contributed to data acquisition, performed most of the data analysis and interpretation, drafted the first draft of the manuscript, critically reviewed the manuscript for important intellectual content.

**Tagreed Mazi** performed part of the data analysis and interpretation, contributed to the first draft of the manuscript, critically reviewed the manuscript for important intellectual content.

**Kari Neier, Noreene M. Shibata, Guillame Jospin, Jonathan A. Eisen, Nathaniel H.O. Harder, Marie C. Heffern, Ashok K. Sharma, Shyam K. More, Maneesh Dave, Shannon M. Schroeder, Li Wang** contributed to data acquisition, contributed with technical and material support, performed part of the data analysis and interpretation and/or statistical analysis, critically reviewed the manuscript for important intellectual content.

**Janine LaSalle, Svetlana Lutsenko** contributed to study concept, provided technical and material support, obtained funding, critical revision of the manuscript for important intellectual content.

**Valentina Medici** designed study concept, obtained funding, contributed to data interpretation, supervised the study, drafted manuscript and provided critical review of the manuscript.

## Data Transparency Statement

The accession number for microbiome-16S rRNA sequencing data is PRJNA909311. Metabolomics and metadata reported in this paper are available via Metabolomics Workbench https://www.metabolomicsworkbench.org/ (ST002423, ST002424, and ST002425).

