## Supplemental Material 1 for "The role of intestine in metabolic dysregulation in murine Wilson disease"

Valentina Medici, M.D., F.A.A.S.L.D.

**Table of Contents**

| Section | Page |
| --- | --- |
| Supplemental Methods | 2 |
| Supplemental Figures | 12 |
| Supplemental Tables | 25 |
| References | 32 |

**Animal models**

All protocols were approved by the UC Davis Institutional Animal Care and Use Committee and followed the National Research Council’s Guide for the Care and Use of Laboratory Animals.

*Atp7b^ΔIEC^ model generation*

*Atp7b*^ΔIEC^ mice were generated by the UC Davis Mouse Biology Program using B6.Cg-Tg(Vil1-cre)997Gum/J mice from the Jackson Laboratory (Bar Harbor, ME) and *Atp7b*^Lox/Lox^ mice^1^ kindly provided by Dr. Lutsenko. Cre-mediated removal of a 1.6-kb fragment in exon 2 results in *Atp7b* inactivation. Vil1-cre mice express Cre recombinase in villus and crypt epithelial cells of the small and large intestine under a villin-1 promoter. A Vil1-cre x *Atp7b*^Lox/Lox^ cross was used to generate mice heterozygous for *Atp7b* Lox and hemizygous for Vil1-cre; Lox^+/-^/Cre^+^ mice were subsequently bred to generate Lox^+/+^/Cre^+^, or *Atp7b*^ΔIEC^, mice. Gene and protein expression was performed to confirm intestinal epithelial cell (IEC)-specific *Atp7b* inactivation (Figure S1). Primer sequences for *Atp7b* were provided by Dr. Lutsenko (Table S1).

*Animal husbandry*

The *Atp7b*^-/-^ global knockout on a C57Bl/6 background (KO) was generated as previously described^1^ and kindly provided by Dr. Svetlana Lutsenko (Johns Hopkins University); C3He-Atp7b^tx-J^/J toxic milk mice (tx-j) and C3HeB/FeJ control mice (C3H) were originally purchased from the Jackson Laboratory. KO, *Atp7b*^ΔIEC^, tx-j, and C3H colonies were bred and maintained on the UC Davis campus in standard plastic shoebox cages with Teklad TEK-Fresh bedding (Envigo, Madison, WI) bedding and nesting enrichment material under the following conditions: 20–23°C, 40%–65% relative humidity, 14 h light/10 h dark light-cycle, and *ad libitum* LabDiet 5001 (PMI, St. Louis, MO) and deionized water. Mice were housed 2-4 per cage. Knockout colonies were maintained by hemizygous (*Atp7b*^ΔIEC^) or heterozygous (KO) breeding to generate littermate controls. KO and *Atp7b*^ΔIEC^ mice were co-housed with their respective littermate controls and segregated by sex at weaning. Tx-j and C3H control were maintained as homozygous colonies. Homozygous tx-j dams have deficient copper levels in their milk for supporting growth and development; tx-j pups must be fostered in a dam with normal milk otherwise they will die from copper deficiency within 14 days post-partum. Tx-j pups were cross-fostered to C3H dams and raised alongside the dam’s biological pups of the same age. Tx-j and C3H mice were

segregated by genotype and sex at weaning.

*Characterization at 16 weeks of age*

At 16 weeks of age, KO and tx-j mice, and their respective controls, had body weights measured then were anesthetized with isoflurane, bled retro-orbitally into K_3_EDTA collection tubes, euthanized by cervical dislocation, and their livers and mesenteric white adipose tissue weighed and flash-frozen in liquid nitrogen. Using disinfected forceps to expel the contents, cecal contents were transferred directly from the cecum into a sterile microfuge tube then flash-frozen in liquid nitrogen. All samples were collected between 9am and 11:30am. Blood samples were centrifuged at 8,000 rpm for 10 minutes and the plasma was aliquoted. All samples were stored at -80°C until further analysis.

*High-fat diet challenge*

From 8 weeks of age, tx-j, KO, and *Atp7b*^ΔIEC^ mice, and their respective controls, were either continued on LabDiet 5001 diet or switched to a 60% kcal fat diet (D12492, Research Diets, Inc., New Brunswick, NJ). After 8 days, mice had body weights measured then were anesthetized with isoflurane, bled retro-orbitally into K_3_EDTA collection tubes, euthanized by cervical dislocation, and the liver weighed and flash-frozen in liquid nitrogen. A small piece of proximal small intestine (~2 cm) was also dissected and flushed with ice-cold Hank’s Balanced Salt Solution for periodate-lysine-paraformaldehyde (PLP) fixation and subsequent immunofluorescent analysis. The intestine piece was placed in 4 ml of PLP fixative (2% paraformaldehyde, 0.0875 M lysine HCl in sodium phosphate buffer, 0.01 M sodium periodate), inverted several times, and fixed at room temperature for 2 hours. The fixative was decanted and the tissue rinsed twice with 0.5 M sucrose in sodium phosphate buffer, 10 minutes each rinse, then held in fresh sucrose solution at 4°C. Blood samples were centrifuged at 8,000 rpm for 10 minutes and the plasma was aliquoted. Plasma and liver samples were stored at -80°C until further analysis.

**IEC isolation**

Mice were anesthetized with isoflurane, then euthanized by cervical dislocation. The ventral incision site was wet with 70% ethanol and the abdominal cavity was opened. The small intestine was cut at the proximal end and gently pulled out of the abdominal cavity to unfurl it, removing the majority of pancreas and mesenteric fat in the process. The small intestine was released by cutting the distal end at the cecum and wet with ice-cold dissection buffer comprised of Hank’s Balanced Salt Solution without calcium/magnesium, 10 mM HEPES, and 5% FBS. Half the full length plus 2 cm of the small intestine was used for dissociation. In a glass petri dish with ice-cold dissection buffer, the intestine was cut into 4-5 smaller segments and each segment was thoroughly flushed with ice-cold dissection buffer. Each segment was then placed on a clean paper towel saturated with ice-cold dissection buffer; any remaining bits of pancreas and mesenteric fat were trimmed; any remaining lumenal content was removed by gently sliding a pair of blunt-tip surgical forceps along the segment (holding the forceps closed), and the segment was held in ice-cold dissection buffer until all segments were cleaned. Each segment was then cut open longitudinally, cut into ~0.5 cm pieces, and placed directly into a conical tube with 40 ml dissociation buffer (HBSS without calcium/magnesium, 10mM HEPES, 12.5 mM EDTA, 5% FBS, 1mM dithiothreitol), and dissociated for 30 minutes at 300 rpm and 37°C in a Corning LSE benchtop orbital shaking incubator (Corning, NY). The dissociated suspension was passed through a 70 µm cell strainer and centrifuged. The supernatant was removed and the cell pellet was washed in 25 ml ice-cold PBS (pH 7.2) with 0.5% FBS then centrifuged. The cells were washed a total of 3 times to remove as many non-IECs, e.g. lymphocytes, as possible. Live cells were counted and the cell suspension was divided across multiple aliquots. Aliquots were centrifuged, supernatant was removed, and cell pellets were flash-frozen and stored at -80°C for downstream analyses. All centrifugations were performed for 5 minutes at 300 x g and 4°C.

**Flow cytometry**

Flow cytometry was performed to determine the quality of the IEC isolation. All antibodies were purchased from Biolegend (San Diego, CA), and dilutions are listed in Table S2. One million live cells were pelleted for 5 minutes at 250 x g and 4°C in 100 µl of PBS then resuspended in 50 µl of staining buffer with EF780 viability dye (Invitrogen, Carlsbad, CA) diluted 1:2000 and incubated for 20 minutes at room temperature in the dark. Cells were then washed with staining buffer, pelleted, and resuspended with 25 µl staining buffer containing anti-mouse CD16/32 antibody to block Fc regions then incubated for 15 minutes at 4°C. A 25 ul antibody cocktail in staining buffer was then added to stain for cell surface markers CD3, CD4, and CD8 (lymphocytes), and Ep-CAM (epithelial cells), and incubated for 30 minutes at 4°C. PE Rat IgG2b k isotype antibody was used as a control. Heat-killed cells stained with EF780 viability dye were used as a control for dead cells. After antibody staining, cells were washed, held in staining buffer, and acquired on a LSRFortessa X-20 (Becton Dickinson, Franklin Lakes, NJ). The data were analyzed using FlowJo software (Ashland, OR). Cells were gated for live cells, excluding the EF780-stained population, with specific marker analyses indicating the successful enrichment of live, viable IECs from *Atp7b*^ΔIEC^ mice.

| **Ep-CAM+ cells** | **CD3+CD4+ T-cells** | **CD3+CD8+ T-cells** |
| --- | --- | --- |
| 87.2 ± 2.3% | 0.004 ± 0.002% | 0.898 ± 0.013% |

**Representative flow cytometric analysis**

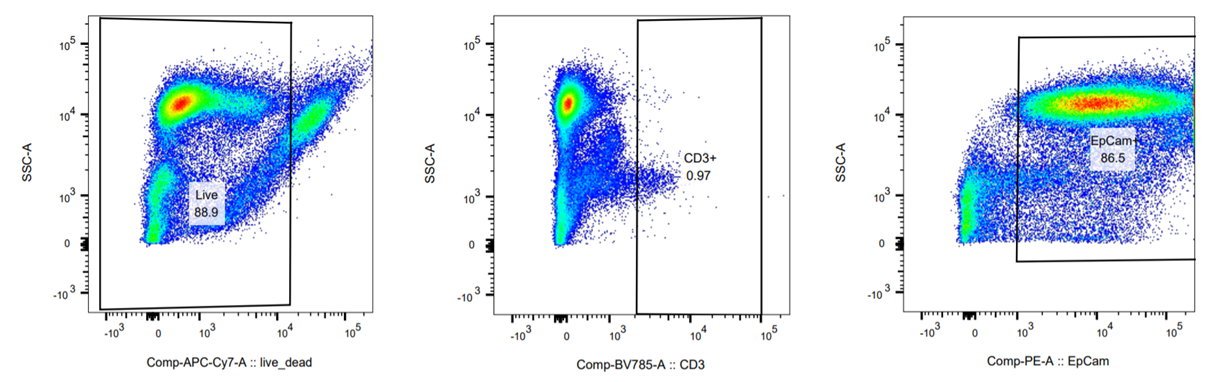

**DNA extraction from cecal contents**

All steps prior to bead lysis were performed in a biosafety cabinet (BSC) cleaned with 70% ethanol and 15 minutes of UV light exposure. Work in the BSC followed standard procedure using 70% ethanol to disinfect gloved hands and all possible equipment and consumables. BSC equipment and consumables set up in advance were also given a 15-minute UV light exposure. DNA extraction was done using the QIAamp Fast DNA Stool Mini Kit (QIAGEN, Valencia, CA) with a modified lysis procedure. Frozen cecal contents were weighed and 200-220 mg were transferred to a sterile 2 ml screw-cap tube containing ~300 mg of 0.1 mm zirconia/silica beads (BioSpec Products, Bartlesville, OK), and 1.6 ml InhibitEX buffer was added. Samples were vortexed until the solution was homogeneous then placed in a Minilys Personal Homogenizer (Bertin Corp, Rockville, MD) and bead-homogenized for 1 minute at 4000 rpm. Samples were chilled on ice for 1 minute then homogenized again for 1 minute at 4000 rpm. Samples were heated at 95°C for 5 minutes with occasional vortexing, given a final vortex for 15 seconds, then centrifuged at 20,000 x g for 5 minutes. Eight hundred µl of supernatant was transferred to a new tube then centrifuged at 20,000 x g for 5 minutes. Six hundred µl of supernatant was transferred to a new tube containing 30 µl proteinase K and 600 µl of Buffer AL was added. After vortexing for 15 seconds, samples were incubated at 70°C for 10 minutes with occasional vortexing. Six hundred µl of 100% ethanol (molecular grade) was added to each sample and thoroughly mixed by vortexing. Six hundred µl of mixture was loaded into a QIAamp spin column, centrifuged at full speed for 1 minute, and the collection tube with flow-through discarded. The column was placed in a new collection tube and the process was repeated 2 more times until all of the sample mixture was passed through the column. All subsequent washes and final elution of total DNA were performed according to the kit protocol. Total DNA concentration and purity were measured with a NanoDrop spectrophotometer (ThermoFisher Scientific, Waltham, MA), and integrity was evaluated by agarose gel electrophoresis.

**16S sequencing analysis**

Library preparation and 16S MiSeq sequencing of cecal content DNA using universal primers for regions V4 and V5 were performed by Integrated Microbiome Resource (Dalhousie University, Halifax, Nova Scotia); 3,116,611 raw PE250 reads were downloaded for processing (39,956.5 average reads per sample, 9,349.6 SD). The reads were de-noised, error-corrected, and merged using the R (3.6.2) and dada2 package version 1.14.1, following the standard online tutorial. Reads were quality controlled by trimming down the end bases to a Q-score of 2 or better. The reverse mates were automatically trimmed down to 180 base pairs due to poor quality. Reads were discarded if they contained Ns (unknown bases) or 2 or more expected errors per mate. Read pairs were merged and de-noised, and 105,763 unique Amplicon Sequence Variants (ASVs) were identified; these ASVs were screened for chimeras (10,849 unique, chimera-free ASVs). After the dada2 pipeline, ASVs were taxonomically annotated using DECIPHER 2.14.0 with default parameters.

The ASV and taxonomy tables were processed using the R package phyloseq (1.30.0). The ASVs were filtered to select for bacteria and remove mitochondria and chloroplasts (4,155 ASVs left). The samples were rarefied to an even depth of 2500 to limit sample exclusion prior to estimating the richness. Bar charts were produced by combining ASVs at various taxonomic rank using the tip_glom function from phyloseq and ampvis2 (2.5.9). The relative abundances in the ASV tables were used to generate the Bray-Curtis dissimilarity matrix and plot the PCoA/NMDS ordination. Beta-diversity analysis was displayed as PCoA plots using permutational multivariate analysis of variance (PERMANOVA). The significantly abundant taxa were identified using the DESeq2 version 1.26.0 R package (Wald test with Benjamini Hochberg correction on parametric variables). Other R packages were used for data manipulation and processing (ape 5.3, tidyverse 1.3.0, dplyr 0.8.5).

MicrobiomeAnalyst (3.6.1) was also used for analyzing and graphical representation of microbiome data.

The prediction of the functional biological pathways in the gut microbiota of the control and WD mouse models was done with a bioinformatics tool, PICRUSt (Phylogenetic Investigation of Communities by Reconstruction of Unobserved States), using a closed-reference OTU table in a biom-format from the script pick_closed_reference_otus.py generated in QIIME. The taxonomy assignment was made with reference sequences from Greengenes database v13.8. The OTU table was then normalized with a PICRUSt workflow using the Langille Lab Online Galaxy Instance (http://galaxy.morganlangille.com) to calculate contributions of various features to known biological pathways based on KEGG orthology groups^2, 3^. MicrobiomeAnalyst (3.6.1)^4, 5^ was used to analyze the PICRUSt-predicted metagenomes to obtain and graphically represent associated pathways between the groups by Classical Univariate Statistical Comparisons analysis by Mann-Whitney/Kruskal-Wallis test.

**Lipidomic profiling**

Untargeted and targeted lipidomic profiling was performed at the UC Davis West Coast Metabolomics Center using charged surface hybrid column/quadrupole time of flight mass spectrometer (LC‐QTOF MS) to detect free fatty acids and complex lipids. Briefly, samples were extracted using methyl tert-butyl ether (MTBE) and water^6^ , and labeled internal standards were added [lysoPE(17:1), lysoPC(17:0), PC(12:0/13:0), PE(17:0/17:0), PG(17:0/17:0), sphingosine (d17:1), d7-cholesterol, SM(17:0), C17 ceramide, d3-palmitic acid, MG(17:0/0:0/0:0), DG(18:1/2:0/0:0), DG(12:0/12:0/0:0), and d5-TG(17:0/17:1/17:0)]. This was followed by ultra-high pressure liquid chromatography on a Waters CSH column, interfaced to a QTOF mass spectrometer operated with electrospray ionization. Data were collected on positive and negative ion mode and processed using MassHunter software (Agilent Technologies, Inc., Santa Clara, CA). Lipids were identified based on MS/MS fragmentation patterns using in-house software, LipidBlast^7^ and lipid quantification was based on internal standards.

For each platform, using known lipids, ion counts in each sample were normalized by scaling counts to the median sum of ion counts for each sample across all mice. Normalized values were log2 transformed. An experiment was defined as a comparison of compound lipid intensities for one biospecimen (liver tissue or plasma) from one platform between two genotypes (C3H vs tx-j, KO vs WT, or *Atp7b*^ΔIEC^ vs iWT). For each experiment, linear regression was used to evaluate intensity differences between genotypes, sex, and their interaction. False discovery rates were calculated for each experiment. Statistical analyses were conducted with R Statistical Computing Software Version 3.6.3.

*Over-representation analysis*

ChemRICH, a statistical enrichment analysis that clusters metabolites based on chemical class using Tanimoto substructure chemical similarity coefficients and calculates cluster significance by Kolmogorov–Smirnov test, was performed and a list of metabolites with corresponding p-values and fold-change were used as input. The outputs were displayed as cluster plots with significantly (p<0.05) impacted clusters of metabolites. Discriminating metabolites based on fold-change and p-value were calculated from normalized metabolite intensities using DESeq package in R. Select metabolites (p<0.05) were generated as a bar plot using ggplot package in R.

**Microbiome and lipidomic correlation analysis**

Compositionally corrected correlation analyses were performed to understand the association between microbial taxa and metabolites using CCREPE function in R. All strongly correlated microbial and metabolite pairs (p<0.05) were used to construct the undirected network using CytoScape version 3.9.0. Nodes in the network plot show microbial taxa and metabolites with color-coded representation of positive (green) and negative (red) correlations.

**RNA extraction and gene expression analysis**

RNA isolation, cDNA generation, and qPCR were performed as previously described^8^. Briefly, RNA was isolated from 25 mg liver and 1x10^7^ IECs using the AllPrep RNA/DNA Mini kit (QIAGEN, Valencia, CA) according to the manufacturer’s instructions. QIAGEN QIAshredder columns were used to homogenize IEC samples; liver samples were homogenized in gentleMACS M-tubes with a gentleMACS Dissociator (Miltenyi Biotec, Auburn, CA) using the RNA_01 program shortened to a 30-second cycle. RNA purity, concentration, and integrity were assessed by NanoDrop spectrophotometer (ThermoFisher Scientific, Waltham, MA) and agarose gel electrophoresis. The Superscript III First-strand Synthesis System (Invitrogen, Carlsbad, CA) was utilized to reverse-transcribe 5 µg RNA into cDNA. Quantitative PCR was run on a ViiA 7 Real-Time PCR System (Applied Biosystems, Foster City, CA) using SYBR Green master mix (Applied Biosystems, Foster City, CA) and a 1/25 cDNA dilution plated in triplicate along with a no-template control. *Gapdh* was used as a reference gene in IECs, *Rplp0* in liver (except for *Ndufs3* in liver for *Atp7b* model validation assay). Statistical significance was determined by Student’s t-test with p<0.05 considered significant.

**Protein extraction and Western blot analysis**

Mouse liver samples (35-40 mg) were homogenized on ice with a hand-held TH homogenizer (OMNI International, Kennesaw, GA) with ice-cold RIPA lysis buffer containing Complete Mini Protease Inhibitor Cocktail and PhosSTOP Phosphatase Inhibitor Cocktail (Roche Diagnostics, Indianapolis, IN). IEC frozen cell pellets (5-17 million cells) were lysed and homogenized in ice-cold RIPA buffer by sonication for 2-5 seconds. Both cell and tissue lysates were centrifuged for 20 minutes at 10,000 rpm and 4°C. The supernatant was collected and protein concentrations were determined with the Pierce BCA Protein Assay Kit (Thermo Fisher Scientific, Waltham, MA) according to the manufacturer’s protocol. All samples and standards were plated in duplicate and read on a Synergy H1 microplate reader (Bio Tek, Winooski, VT).

For ATP7B: Equal amounts of protein (50 µg) were denatured in 4X LDS sample buffer (Thermo Fisher Scientific, Waltham, MA) with 2% Beta-mercaptoethanol (BIO-RAD, Hercules, CA). For iWT and *Atp7b*^ΔIEC^ IEC samples, urea was added for a final urea concentration of 1.33 M in each sample. Proteins were separated by SDS-PAGE and transferred onto a pre-activated PVDF membrane using the BIO-RAD Trans-Blot Turbo transfer system. Membranes were blocked with 5% protease and fatty acid-free BSA (MilliporeSigma, Burlington, MA) in tris-buffered saline with Tween-20 (TBST) for 1 hour then probed with ATP7B or β-ACTIN primary antibody overnight at 4°C. Membranes were washed in TBST then incubated with the appropriate horseradish peroxidase-conjugated anti-rabbit (ATP7B) and Alexafluor 555 conjugated anti-mouse IgG secondary antibody (β-ACTIN) for 1 hour at room temperature. Membranes were washed in TBST again and blots were developed with Clarity Western ECL Substrate (Thermo Fisher Scientific, Waltham, MA). Membranes were visualized using a Chemidoc MD system (BIO-RAD, Hercules, CA).

For all other proteins: Equal amounts of protein (25 µg) were denatured 1:1 in 2X Laemmli buffer (BIO-RAD, Hercules, CA) at 95°C for 10 minutes then held on ice. Proteins were separated by SDS-PAGE and transferred onto a pre-activated PVDF membrane using the BIO-RAD Trans-Blot Turbo transfer system. Membranes were blocked with 5% non-fat milk in TBST for 2 hours then probed with the desired primary antibodies overnight at 4°C. Membranes were washed in TBST then incubated with the appropriate horseradish peroxidase-conjugated anti-rabbit or anti-mouse IgG secondary antibody for 1 hour at room temperature. Membranes were washed in TBST again and blots were developed with Clarity Western ECL Substrate. Membranes were visualized using a FujiFilm LAS-4000 imaging system. Densitometry analyses were quantified by using Fujifilm Multi Gauge software and normalized to β-ACTIN. Statistical significance was determined by Student’s t-test with p<0.05 considered significant. Antibodies and dilutions are listed below in Table S2**.**

**ApoB48 immunofluorescence**

PLP-fixed liver was transferred from sucrose solution to a cryomold for O.C.T. compound embedding. O.C.T.-embedded tissues were frozen and stored at -80°C until sectioning and slide mounting. Slides were stored at -80°C. For staining, the slide box was removed from -80ºC and allowed to sit at room temperature for 20 minutes. Slides were washed 4 times with PBS, 5 minutes per wash, to completely remove the O.C.T. compound. Slides were immersed in blocking solution (1% BSA + 0.1% Tween 20 in PBS) at room temperature for 30 minutes. Slides were then incubated with the anti-ApoB primary antibody in a humidity chamber at 4ºC overnight. Slides were washed 4 times with 0.1% Tween 20 in PBS (PBST), 5 minutes per wash. Alexa Fluor™ Plus 647 Phalloidin (Invitrogen, Carlsbad, CA) was used to stain F-actin and prepared according to manufacturer's instructions then added to the secondary antibody solution. Secondary antibodies were added onto each slide and incubated at room temperature for 45 minutes. Slides were washed twice with PBST for 5 minutes each, and then once with PBS for 5 minutes. ProLong™ Gold Antifade Mountant with DAPI (Invitrogen, Carlsbad, CA) was applied to the slides. Slides were imaged with a Zeiss Confocal Microscope LSM800.

**H&E and Oil Red O Histology**

Formalin-fixed liver was paraffin-embedded, sectioned, mounted, stained, and imaged, and O.C.T. compound-embedded liver was sectioned, mounted, stained, and imaged by the UC Davis Center for Genomic Pathology Laboratory.

**Figure S1. *Atp7b*^ΔIEC^ model validation.**

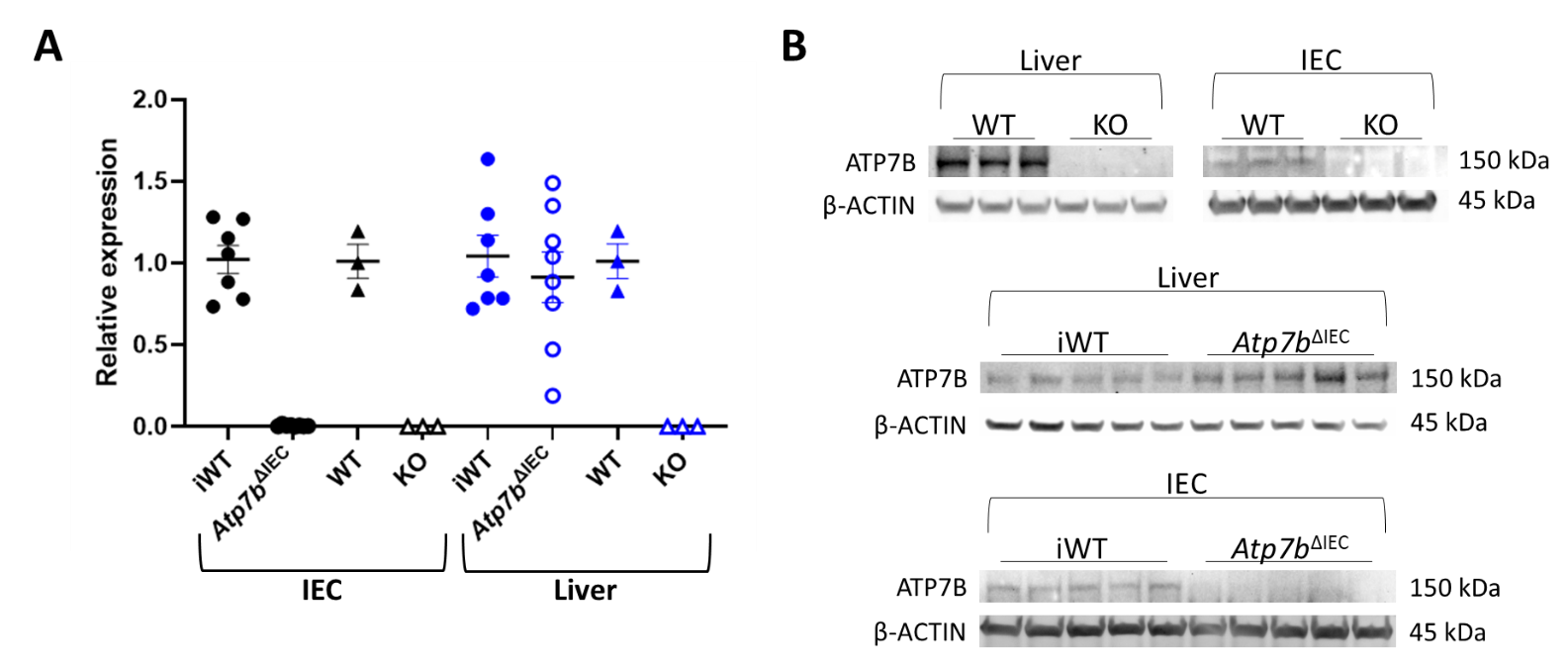

A: Quantitative PCR of *Atp7b* was performed for 9-week-old *Atp7b*^ΔIEC^ (Lox^+/+^/Cre^+^, n=8) mice and their respective littermate control iWT (Lox^+/+^/Cre^-^, n=7). KO mice (n=3) and their respective littermate control WT (n=3) were also included as additional assay controls. Results were normalized to *Gapdh* in IECs and *Ndufs3* in liver. *Atp7b* expression is reduced 99% in *Atp7b*^ΔIEC^ Cre^+^ IECs compared to Cre^-^. This dramatic, but incomplete, reduction of *Atp7b* expression in the intestine-specific Cre-lox mouse model is commensurate with expression levels demonstrated by Muchenditsi et al^1^ in the hepatocyte-specific Cre-lox mouse model, *Atp7b*^ΔHep^. Residual expression detected may be due to either contaminating lymphocytes or incomplete inactivation by Cre recombinase in a hemizygous state. KO mice demonstrated no detectable *Atp7b* transcript levels. *Atp7b* levels were not different between *Atp7b*^ΔIEC^ and iWT samples in liver, indicating IEC specificity. B: Representative ATP7B immunoblot of 9-week-old KO (n=3) and *Atp7b*^ΔIEC^ (n=5) liver and IEC lysates and their respective littermate controls. Blots were normalized to β-ACTIN. ATP7B protein expression reflects gene expression in specificity and quantity; ATP7B is not detectable in KO mice but, although dramatically reduced, ATP7B can be detected in *Atp7b*^ΔIEC^ mice as a very faint band.

*Atp7b*, ATPase copper transporting beta; *Atp7b*^ΔIEC^ mice, intestine-specific *Atp7b* knockout mice; *Gapdh*, glyceraldehyde 3-phosphate dehydrogenase; IECs, intestinal epithelial cells; iWT, littermate controls (Lox^+/+^/Cre^-^) for *Atp7b*^ΔIEC^mice; KO, *Atp7b* null global knockout mice on a C57Bl/6 background; *Ndufs3*, NADH:ubiquinone oxidoreductase core subunit S3; WT, littermate controls (*Atp7b*^+/+^) for KO mice.

**Figure S2. Study design**

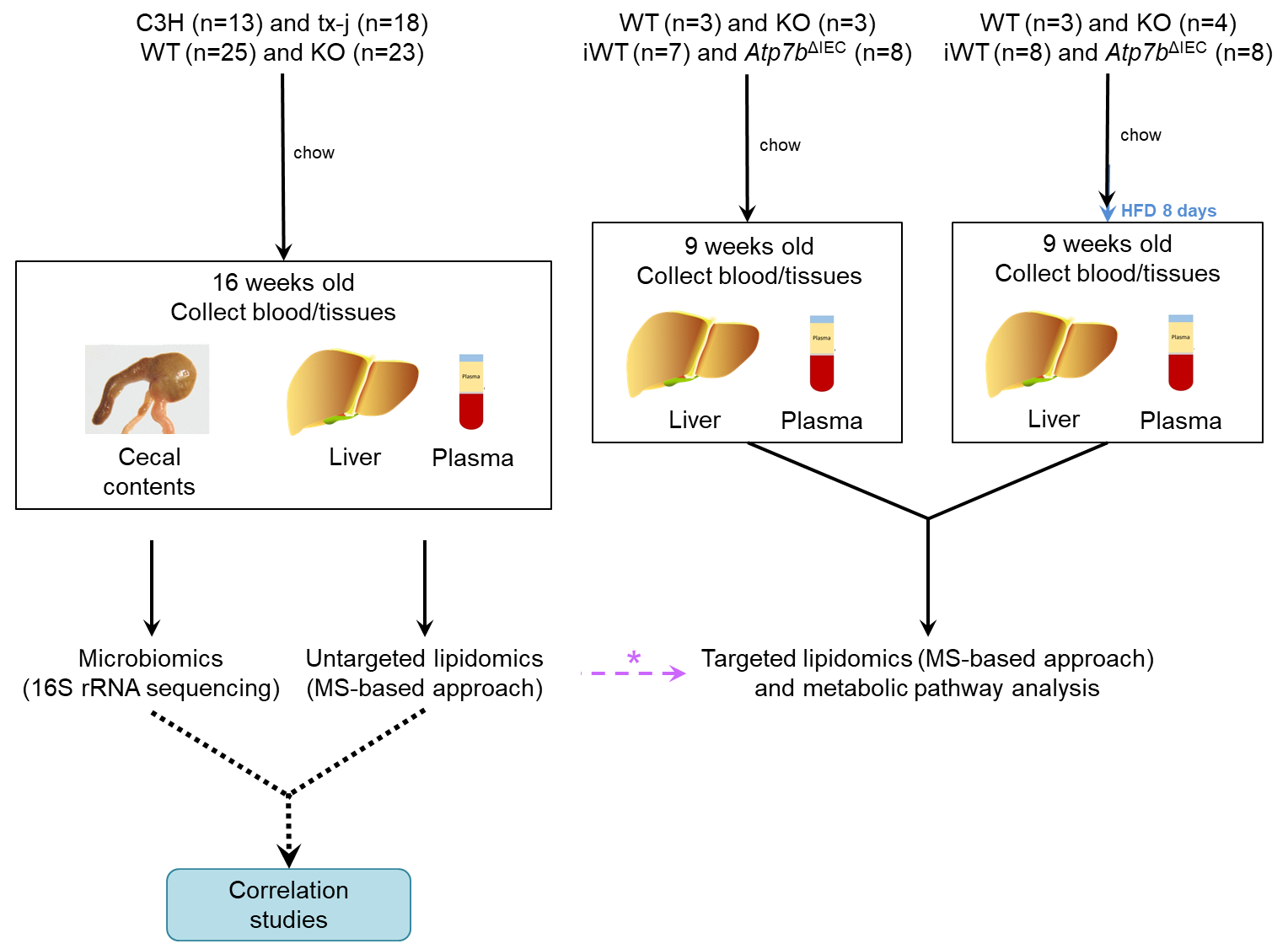

*The most differentially abundant lipid classes from the 16-week untargeted analysis were selected for targeted lipidomics in both 9-week liver and plasma. Chow and HFD were investigated in both KO and *Atp7b*^ΔIEC^ mice to determine metabolic effects of global *Atp7b* deficiency as well as intestine-specific metabolic effects without the presence of hepatic copper accumulation and liver disease.

*Atp7b*^ΔIEC^, intestine-specific *Atp7b* knockout mice; C3H, Jackson Laboratory C3HeB/FeJ control for tx-j, chow; Purina LabDiet 5001; HFD, high-fat diet Research Diets D12492; iWT, littermate controls (Lox^+/+^/Cre^-^) for *Atp7b*^ΔIEC^ mice; KO, *Atp7b* null global knockout mice on a C57Bl/6 background; MS, mass spectrometry; tx-j, Jackson Laboratory toxic-milk mouse C3He-Atp7b^tx-J^/J; WT, littermate controls (*Atp7b*^+/+^) for KO mice.

**Figure S3. Tx-j and KO mice liver histology at 16 weeks of age.**

**1a**

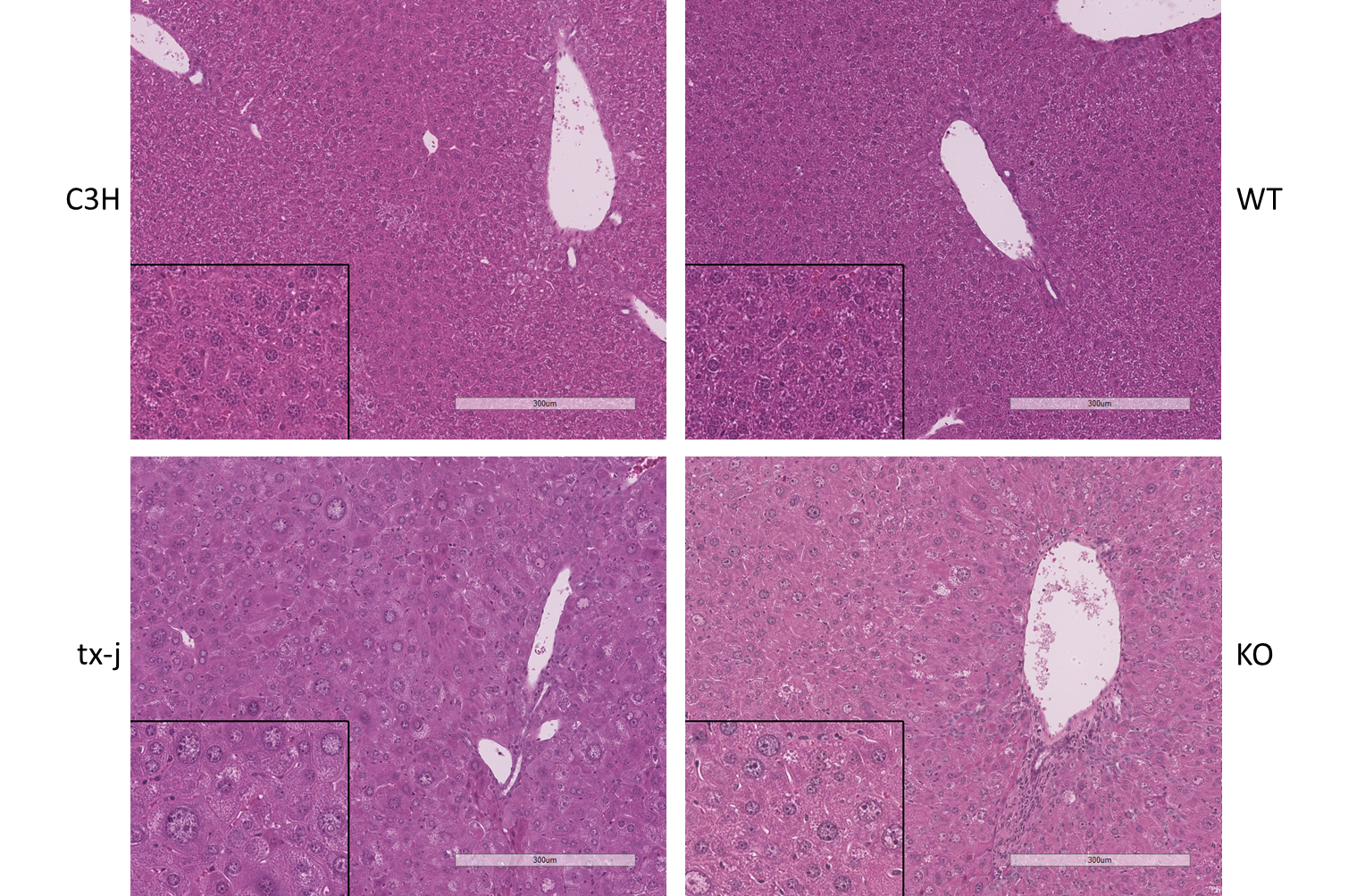

**2b**

**2a**

**1b**

Representative hematoxilin and eosin staining in mouse livers at 16 weeks of age on Purina LabDiet 5001. 10X magnification with 20X insert. Scale bar = 300 µm at 10X.

Liver histology is normal in control C3H (1a) and WT (1b) mice. Tx-j (2a) and KO (2b) mice presented diffusely enlarged and glycogenated hepatocyte nuclei, and minimal cytosolic glycogenosis.

C3H, C3HeB/FeJ controls for tx-j; KO, *Atp7b* null global knockout mice on a C57Bl/6 background; tx-j, Jackson Laboratory toxic-milk mouse C3He-Atp7b^tx-J^/J; WT, littermate controls (*Atp7b*^+/+^) for KO mice.

**Figure S4. Taxonomic profile for tx-j vs C3H mice.**

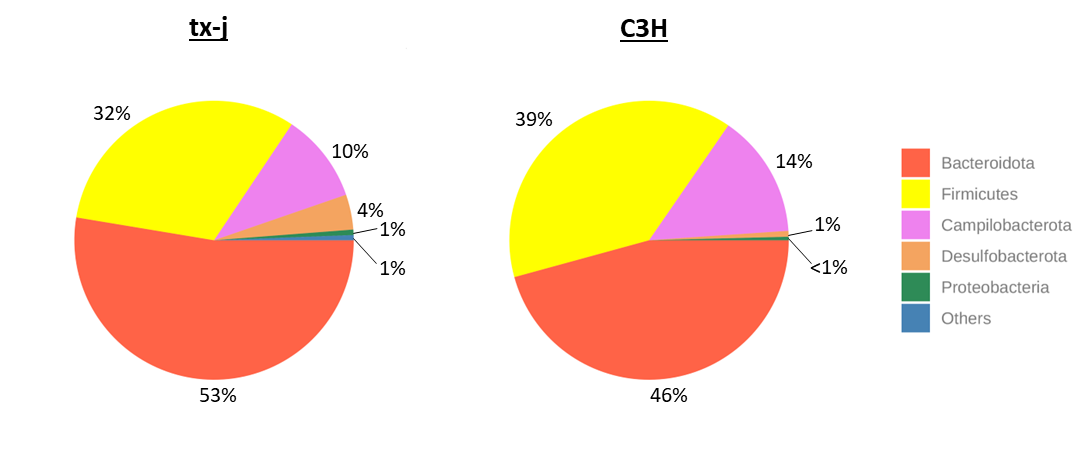

Pie chart representation of the relative abundance of relevant bacterial phyla in tx-j vs C3H mice. Phyla are identified by colors, as indicated in the legend, and numbers are the relative abundance in percentage.

C3H, Jackson Laboratory C3HeB/FeJ control for tx-j; tx-j, Jackson Laboratory toxic-milk mouse C3He-Atp7b^tx-J^/J.

**Figure S5. Taxonomic profile for KO vs WT mice.**

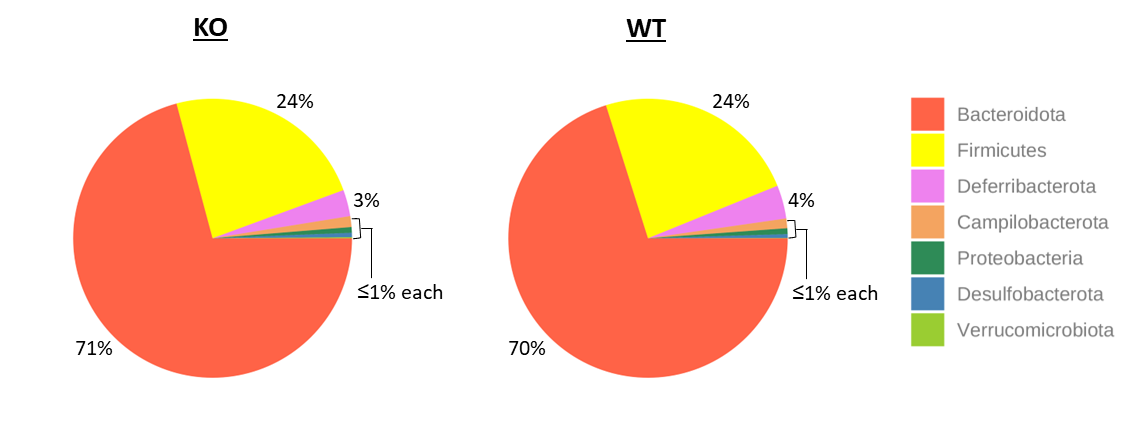

Pie chart representation of the relative abundance of relevant bacterial phyla in KO vs WT mice. Phyla are identified by colors, as indicated in the legend, and numbers are the relative abundance in percentage.

KO, *Atp7b* null global knockout mice on a C57Bl/6 background; WT, littermate controls (*Atp7b*^+/+^) for KO mice.

**Figure S6. Variation of taxonomic abundance at the genus level by genotype and sex in tx-j vs C3H mice.**

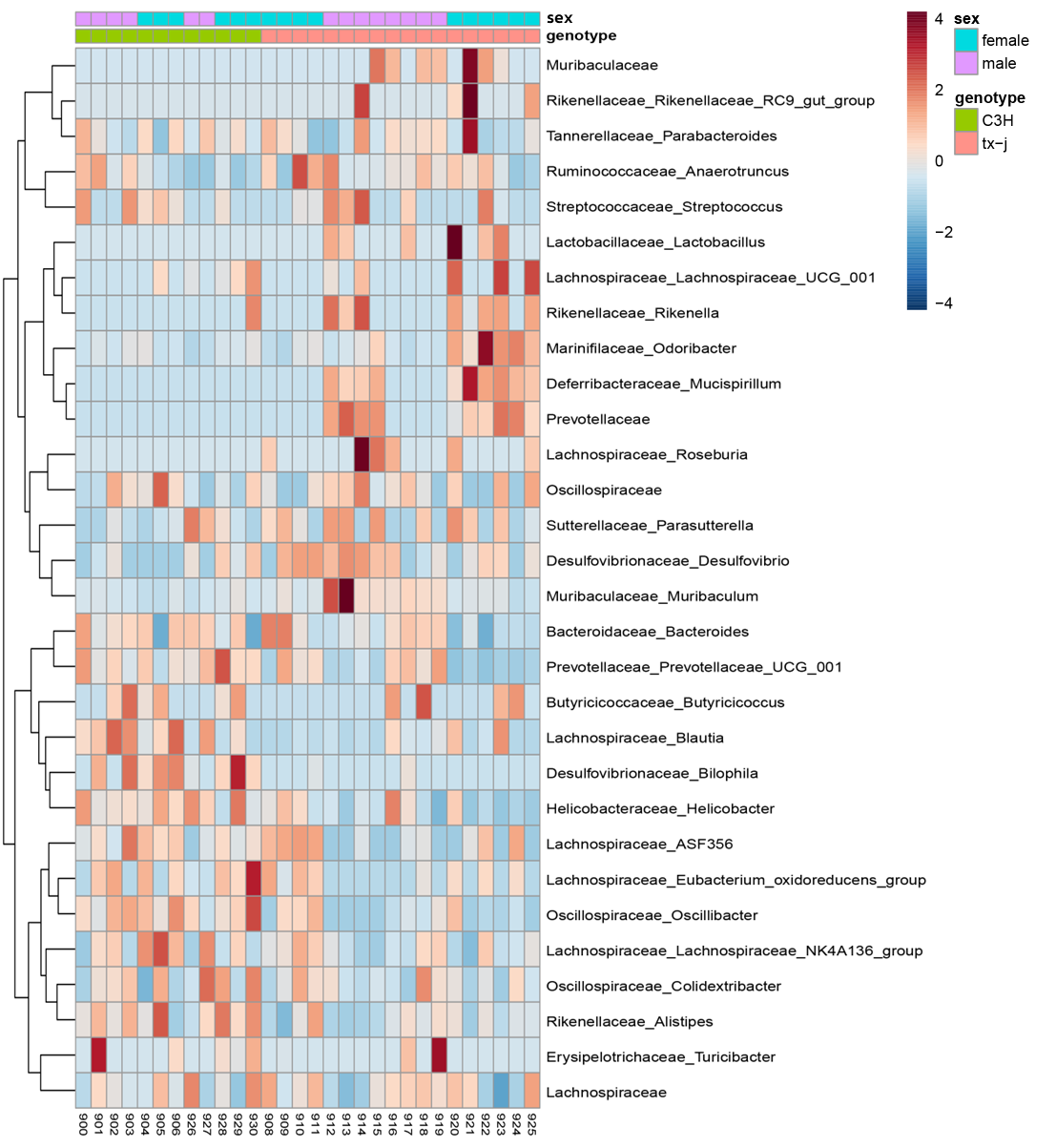

Hierarchical heat map clustering of the samples based on Pearson’s correlation coefficient for the abundance of family- and genus-level taxa. The color scale indicates the scaled abundance of each variable, denoted as the Z-score: red, high abundance; blue, low abundance. Column bars are colored according to genotype (green, C3H; red, tx-j) and sex (turquoise, male; lavender, female).

C3H, C3HeB/FeJ control for tx-j; tx-j, Jackson Laboratory toxic-milk mouse C3He-Atp7b^tx-J^/J.

**Figure S7. Principal component analysis of tx-j vs C3H mice.**

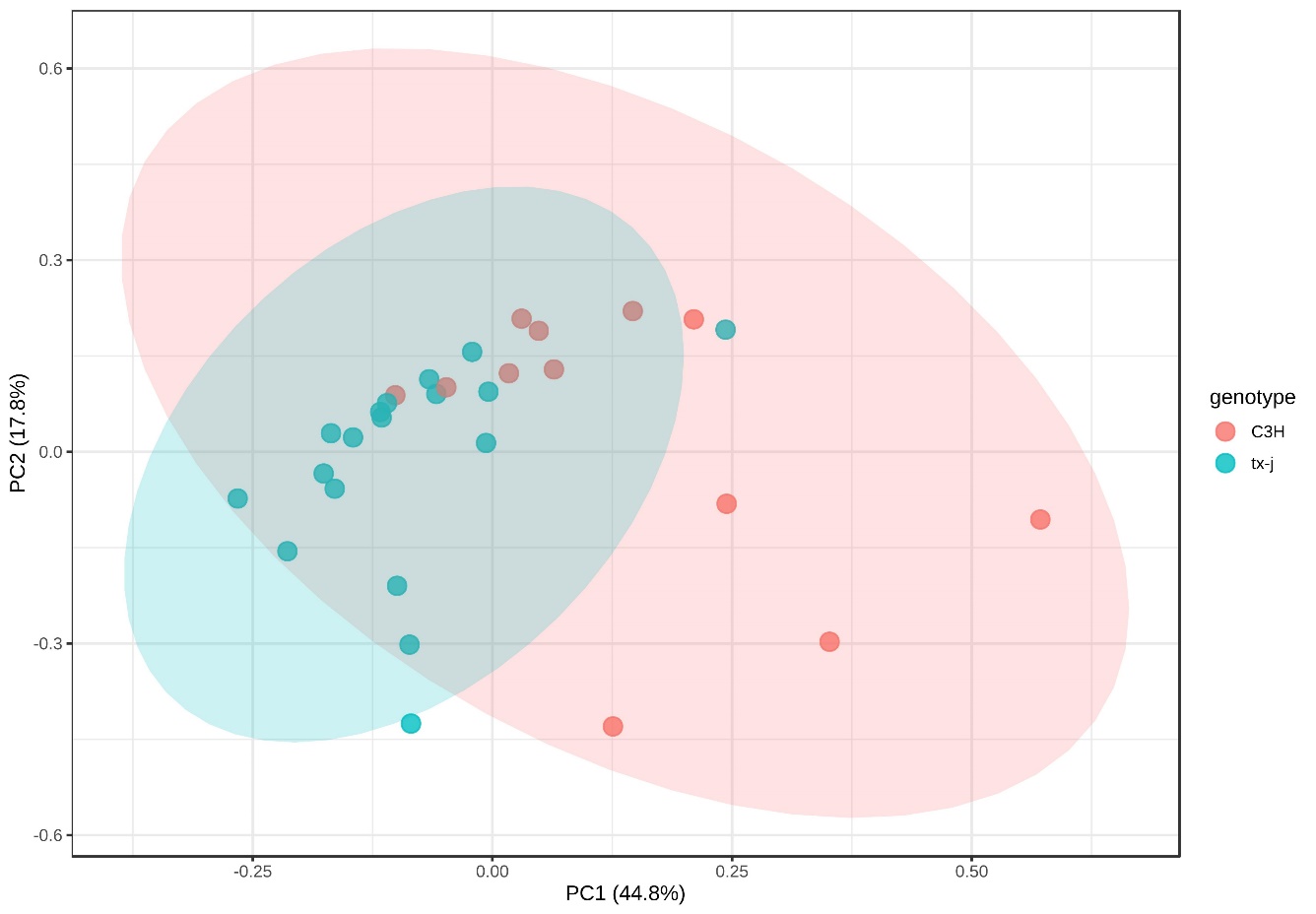

Principal component analysis based on the KEGG ortholog abundances profile of the gut microbiota in tx-j vs C3H mice with 44.8% of variance between the groups.

C3H, Jackson Laboratory C3HeB/FeJ control for tx-j; tx-j, Jackson Laboratory toxic-milk mouse C3He-Atp7b^tx-J^/J.

**Figure S8. Liver lipid modules associated with tx-j and KO mice.**

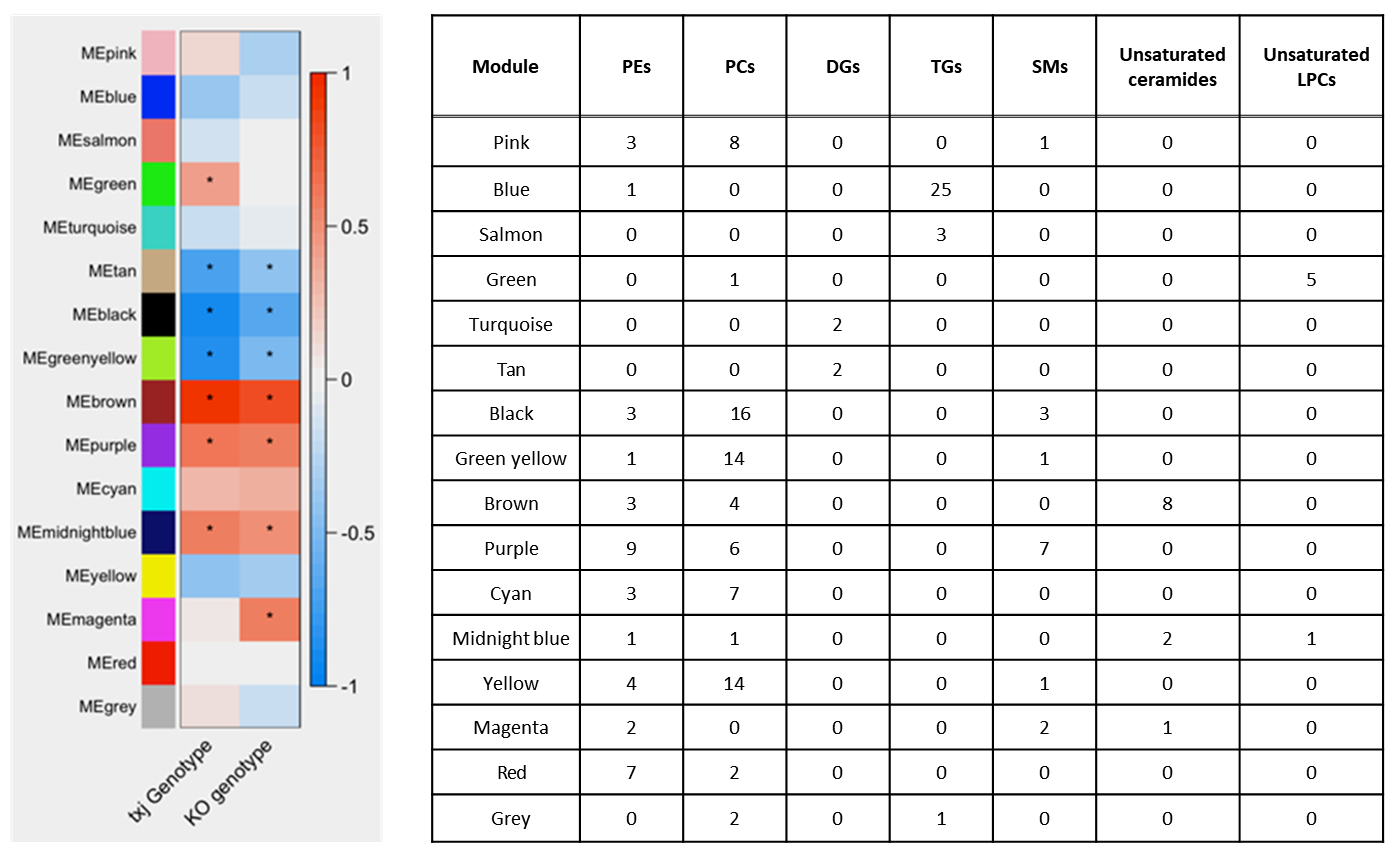

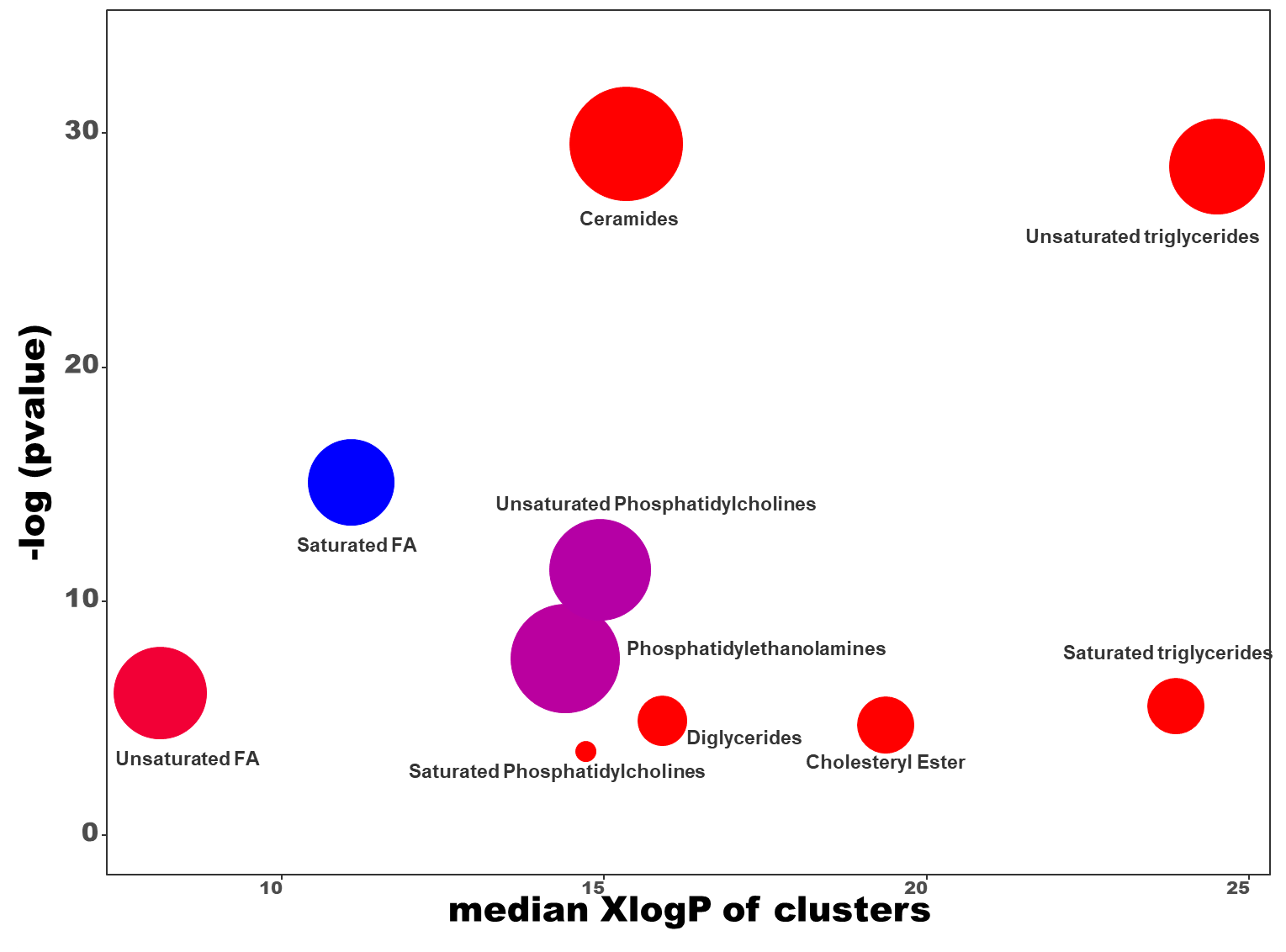

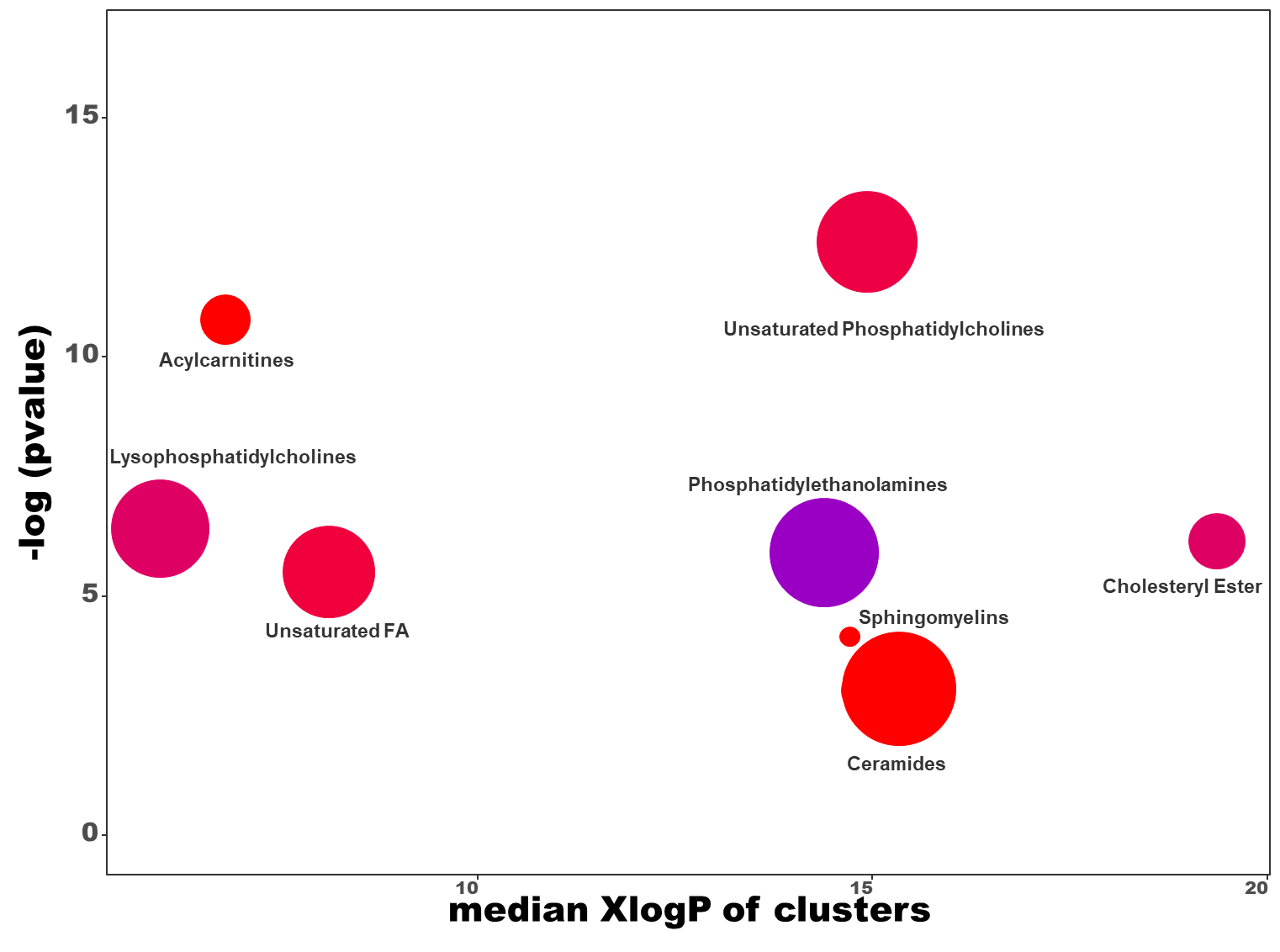

**F**

**E**

**Plasma**

**Liver**

Correlation between the microbiome and lipids using a weighted co-network analysis and color-coded modules. Correlation coefficients of 16 liver lipid modules are represented in the heat map. Red and blue heat map squares indicate positive and negative associations, respectively. An asterisk (*) indicates a FDR-corrected p-value ≤ 0.05. Values in the color legend represent the number of lipids in each class within a color-coded module.

DGs, diglycerides; LPCs, lysophosphatidylcholines; PCs, phosphatidylcholines; PEs, phosphatidylethanolamines; SMs, sphingomyelins; TGs, triglycerides.

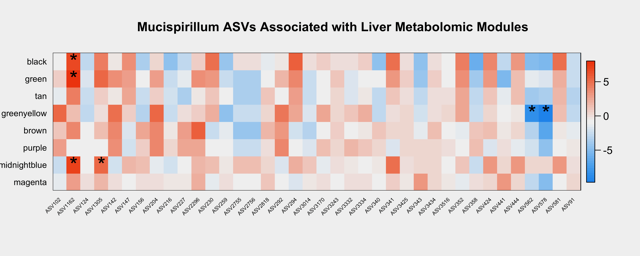
**Figure S9. *Mucispirillum* amplicon sequence variants associated with liver lipid modules**.

Correlation between *Mucispirillum* amplicon sequence variants and liver lipid modules using weighted co-network analysis. Correlation coefficients of 8 liver lipid modules are represented in the heat map. Red and blue heat map squares indicate positive and negative associations, respectively. An asterisk (*) indicates a FDR-corrected p-value ≤ 0.05. See table in Figure S8 for module color legend.

**Figure S10. *Atp7b*^ΔIEC^ liver histology at 9 weeks of age.**

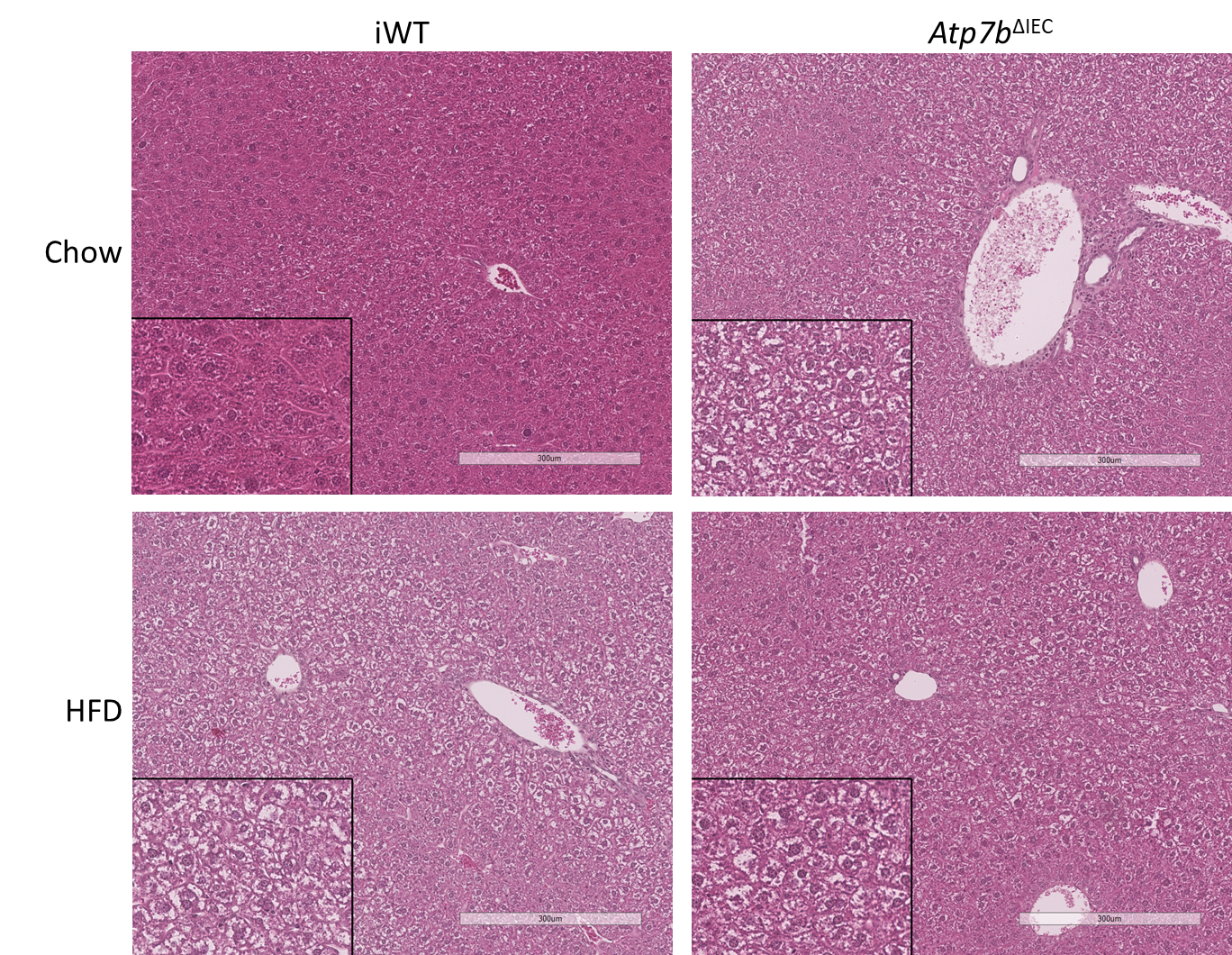

**1a**

**1b**

**2b**

**2a**

Representative hematoxilin and eosin staining in mouse livers at 9 weeks of age on Purina LabDiet 5001 (1a, 1b) and after an 8-day HFD challenge (2a, 2b). 10X magnification with 20X insert. Scale bar = 300 µm at 10X.

Liver histology is normal in control iWT (1a) mice. *Atp7b*^ΔIEC^ (1b) presented marked cytosolic glycogenosis and vacuolization with mild steatosis, similar to the findings observed in iWT mice on HFD (2a) and *Atp7b*^ΔIEC^ on HFD (2b).

*Atp7b*^ΔIEC^, intestine-specific *Atp7b* knockout mice; HFD, high-fat diet Research Diets D12492; iWT, littermate controls (Lox^+/+^/Cre^-^) for *Atp7b*^ΔIEC^ mice.

**Figure S11. Lipidomic profile in liver and plasma of *Atp7b*^ΔIEC^ compared with iWT control.**

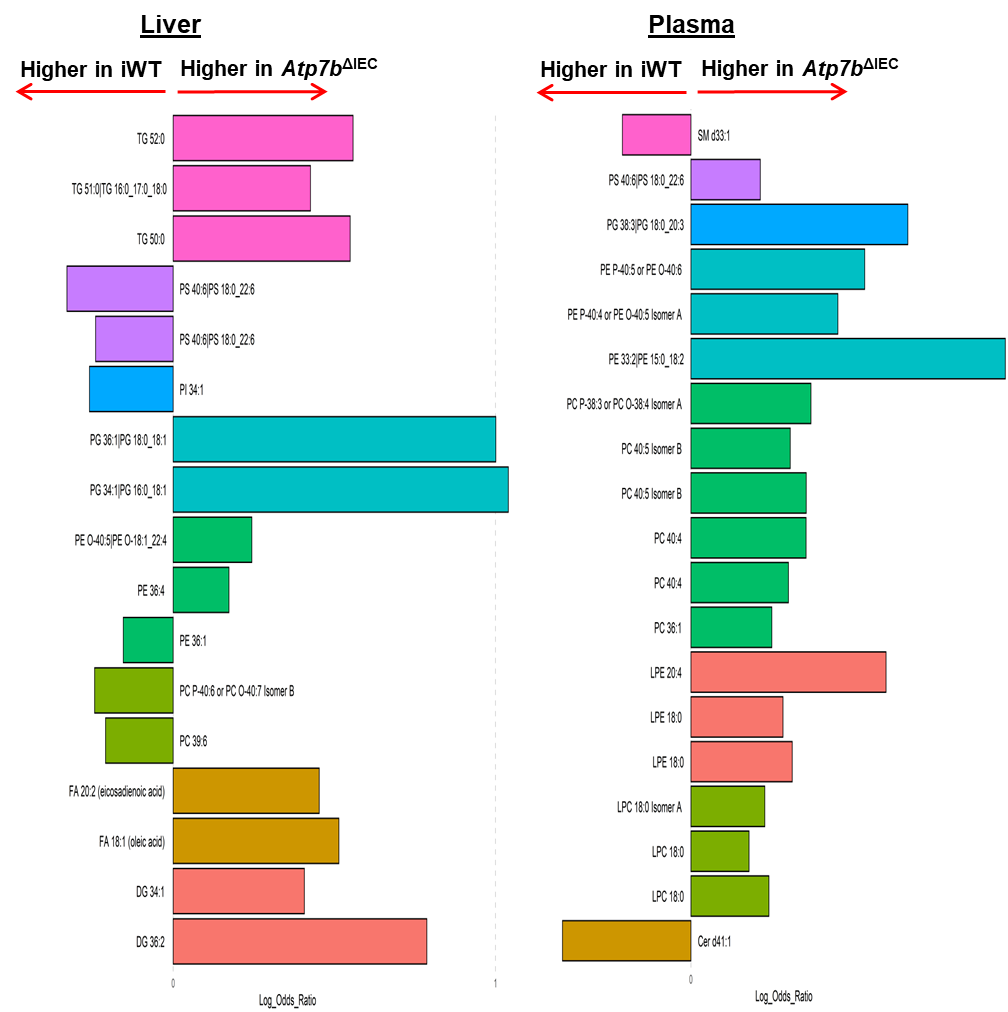

The LEfSe plot represents a logarithm of the average level ratios of statistically different lipids between *Atp7b*^ΔIEC^ and WT mice in liver (A) and plasma (B). A positive log ratio bar indicates higher lipid levels in *Atp7b*^ΔIEC^ mice; a negative log ratio bar indicates higher levels in iWT mice. Colors represent lipids in the same

class.

*Atp7b*^ΔIEC^, intestine-specific *Atp7b* knockout mice; iWT, littermate controls (Lox^+/+^/Cre^-^) for *Atp7b*^ΔIEC^ mice; LEfSe, linear discriminant analysis effect size.

**Figure S12:** **Lipidomic profiles in liver and plasma of KO mice fed chow or HFD.**

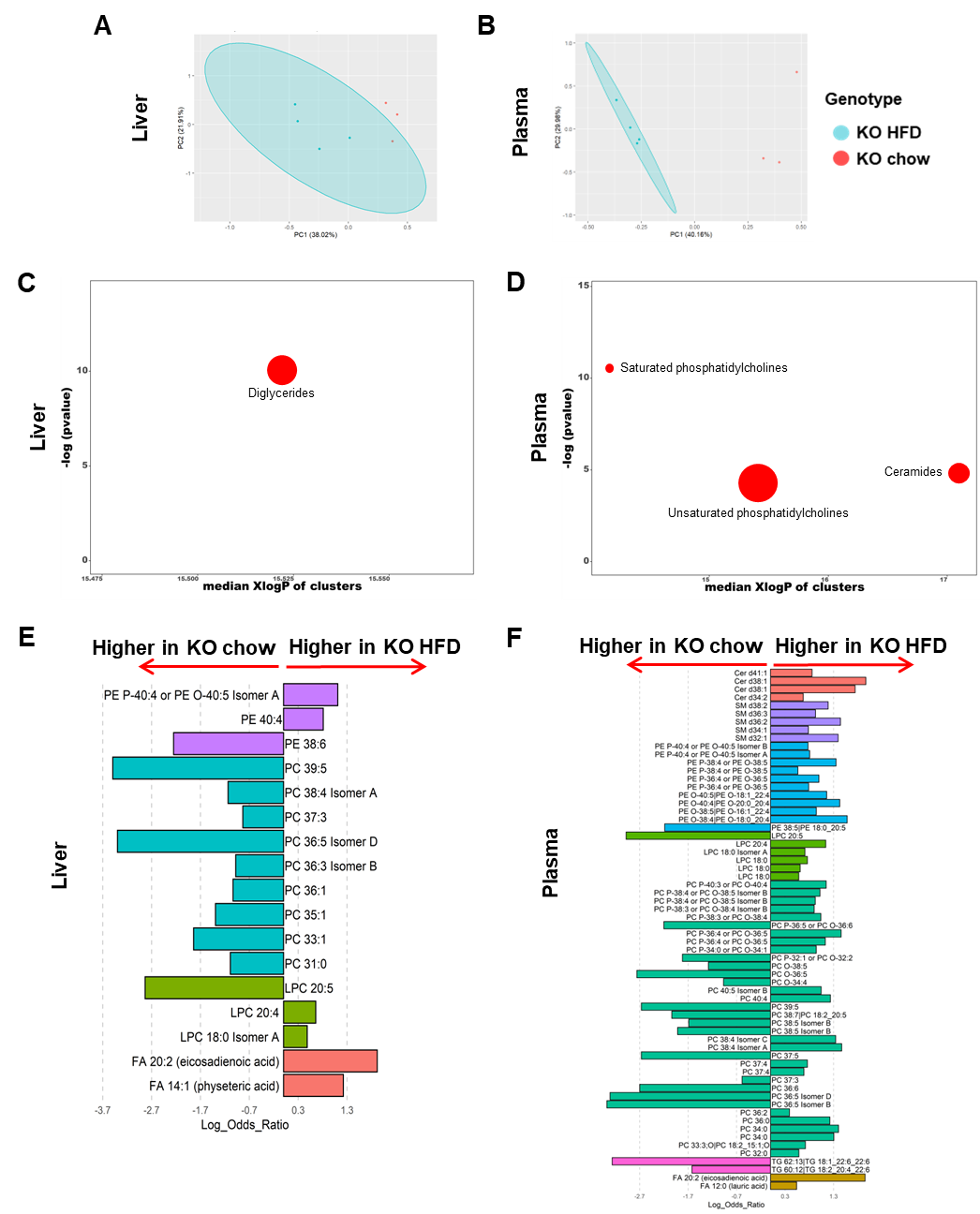

A, B: Principal component analysis based on the lipid profiling of KO mice fed chow or HFD in the liver and plasma. C, D: Chemical similarity enrichment analysis (ChemRICH) and enrichment statistics plot for KO HFD vs KO chow mice in liver and plasma. Each cluster represents an altered chemical class of metabolites (p<0.05). Cluster size represents the total number of metabolites. Cluster color represents the directionality of metabolite differences: red, higher in KO HFD mice. The x-axis represents the cluster order on the chemical similarity tree. The y-axis represents chemical enrichment p-values calculated using a Kolmogorov–Smirnov test. E, F: Linear discriminant analysis effect size (LEfSe) plots representing the logarithm of the ratios of the average levels of statistically different lipids between KO HFD and KO chow mice in liver and plasma. A positive log ratio bar indicates higher lipid levels in KO HFD; a negative log ratio bar indicates higher levels in KO chow mice. Colors represent lipids in the same class.

**Table S1. Primer sequences for gene expression analysis.**

| *Gene* |  |  | Primers 5’ – 3’ | Exon spanning | Amplicon  (bp) | % efficiency |
| --- | --- | --- | --- | --- | --- | --- |
| *Atp7b** | ATPase copper transporting beta | F | TTCTTGCGTCAAGTCCATTG | No | 158 | 102.7 |
|  |  | R | CCTCAAAGCCCATATCCTCA | No |  |  |
|  |  | R | CCGTGGGTGATCGTGTCTGG | No |  |  |
| *Scd1* | stearoyl-Coenzyme A desaturase 1 | F | TCATTCTCATGGTCCTGCTGC | No | 113 | 95.3 |
|  |  | R | CAGAGCGCTGGTCATGTAGTAGA | No |  |  |
|  |  | R | AGGACTCCAAAGGACAGGCTCT | No |  |  |

*Primers are against exon 2 which is where the loxP modification site is located.

**Table S2. Antibodies and dilutions.**

| **Procedure** | **Antibody** | **Dilution** | **Vendor** | **catalog number** |
| --- | --- | --- | --- | --- |
| Flow cytometry | anti-mouse CD16/32 | 1:32 | Biolegend  (San Diego, CA) | 101302 |
| Flow cytometry | Brilliant Violet 711 anti-mouse CD4 | 1:200 | Biolegend  (San Diego, CA) | 100550 |
| Flow cytometry | Brilliant Violet 605 anti-mouse CD8a | 1:100 | Biolegend  (San Diego, CA) | 100744 |
| Flow cytometry | Brilliant Violet 785 anti-mouse CD3 | 1:40 | Biolegend  (San Diego, CA) | 100232 |
| Flow cytometry | PE anti-mouse CD326 (Ep-CAM) | 1:80 | Biolegend  (San Diego, CA) | 118206 |
| Flow cytometry | PE Rat IgG2b k isotype | 1:80 | Biolegend  (San Diego, CA) | 400636 |
| Western blot | AMPKα | 1:1000 | Cell Signaling Technology  (Danvers, MA) | 23A3 |
| Western blot | ATP7B | 1:1000 | Abcam  (Burlingame, CA) | ab124973 |
| Western blot | β-ACTIN (for ATP7B) | 1:3000 | Cell Signaling Technology (Danvers, MA) | 4970S |
| Western blot | β-ACTIN | 1:5000 | Sigma-Aldrich  (St. Louis, MO) | A5441 |
| Western blot | pAMPKα | 1:1000 | Cell Signaling Technology (Danvers, MA) | 2535 |
| Western blot | PPARα | 1:500 | Abcam  (Burlingame, CA) | ab24509 |
| Western blot | PPARγ | 1:1000 | Cell Signaling Technology (Danvers, MA) | 2443S |
| Western blot | Anti-mouse secondary | 1:10000 | Jackson Immunoresearch Lab  (West Grove, PA) | 115-035-003 |
| Western blot | Anti-rabbit secondary | 1:10000 | Jackson Immunoresearch Lab  (West Grove, PA) | 111-035003 |
| Western blot | Anti-rabbit secondary (for ATP7B) | 1:2000 | Cell Signaling Technology (Danvers, MA) | 7074S |
| Western blot | Anti-mouse secondary (for ATP7B) | 1:5000 | Thermo Fisher Scientific Waltham, MA) | A28180 |

**Table S3. Gross phenotypes of mouse models at 16 weeks of age.**

|  | **n** | | **Body weight (g)** | | **Liver / Body weight** | | **MWAT / Body weight** | |
| --- | --- | --- | --- | --- | --- | --- | --- | --- |
|  | M | F | Male | Female | Male | Female | Male | Female |
| **C3H** | 6 | 6 | 32.6 ± 1.3 | 30.1 ± 2.0^*^ | 0.053 ± 0.003 | 0.055 ± 0.003 | 0.0074 ± 0.0011 | 0.0101 ± 0.0033 |
| **tx-j** | 10 | 8 | 26.9 ± 1.1**^††^** | 22.8 ± 0.7^**^**^††^** | 0.054 ± 0.003 | 0.060 ± 0.002^*^**^††^** | 0.0059 ± 0.0012**^†^** | 0.0049 ± 0.0012**^†^** |
| **WT** | 12 | 13 | 29.2 ± 1.8 | 23.6 ± 1.7^**^ | 0.049 ± 0.006 | 0.045 ± 0.007 | 0.0042 ± 0.0012 | 0.0047 ± 0.0025 |
| **KO** | 11 | 11 | 27.7 ± 1.7 | 23.0 ± 1.1^**^ | 0.053 ± 0.003 | 0.056 ± 0.011**^†^** | 0.0036 ± 0.0008 | 0.0036 ± 0.0010 |

Values are means ± SD and statistical significance was determined by Student’s t-test. An asterisk (*) indicates values are significantly different between sexes within the same genotype (* p<0.01, ** p<0.001). A dagger (**†**) indicates values are significantly different between a WD model (tx-j or KO) and its respective control (C3H or WT) within the same sex (**^†^** p<0.05, **^††^** p<0.001).

C3H, Jackson Laboratory C3HeB/FeJ controls for tx-j; KO , *Atp7b* null global knockout on C57Bl/6 background; MWAT, mesenteric white adipose tissue; tx-j, Jackson Laboratory toxic milk mouse C3He-Atp7b^tx-J^/J; WT, littermate controls (*Atp7b*^+/+^) for KO.

**Table S4. Bacterial genus abundance for tx-j vs C3H.**

| **Genus** | **log2FC** | **LEfSe** | **p-value** |
| --- | --- | --- | --- |
| *Mucispirillum* | 10.417 | 1.751 | 2.70E-09 |
| *Lactobacillus* | 3.8705 | 0.99431 | 9.92E-05 |
| *Muribaculum* | 7.099 | 1.8679 | 0.000144 |
| *Helicobacter* | -0.60362 | 0.1953 | 0.001996 |
| *Oscillibacter* | -1.4366 | 0.46987 | 0.002233 |
| *Bilophila* | -2.7472 | 0.95068 | 0.003856 |
| *Odoribacter* | 1.521 | 0.53052 | 0.004143 |
| *Rikenellaceae_RC9_gut_group* | 1.5359 | 0.55945 | 0.006044 |
| *Blautia* | -2.0482 | 0.79176 | 0.009683 |
| *Alistipes* | -0.65048 | 0.3028 | 0.031698 |
| *Colidextribacter* | -0.42969 | 0.23987 | 0.073241 |
| *Lachnospiraceae_NK4A136_group* | -0.63345 | 0.37261 | 0.08912 |
| *Prevotellaceae_UCG_001* | -0.87204 | 0.54265 | 0.10805 |
| *Rikenella* | 2.089 | 1.5711 | 0.18365 |
| *Anaerotruncus* | 0.82786 | 0.62536 | 0.18557 |
| *Desulfovibrio* | 0.69534 | 0.6335 | 0.27237 |
| *Turicibacter* | -1.9271 | 1.7571 | 0.27274 |
| *Roseburia* | 1.106 | 1.0303 | 0.28304 |
| *Eubacterium_oxidoreducens_group* | -1.0068 | 1.2095 | 0.40519 |
| *Parasutterella* | 0.40986 | 0.55204 | 0.45781 |
| *Bacteroides* | -0.17254 | 0.28569 | 0.54589 |
| *Parabacteroides* | 0.20447 | 0.35882 | 0.5688 |
| *Butyricicoccus* | -0.65095 | 1.1819 | 0.58181 |
| *Lachnospiraceae_UCG_001* | -0.71723 | 1.5492 | 0.64339 |
| *Streptococcus* | 0.3348 | 0.85022 | 0.69375 |
| *ASF356* | -0.07047 | 0.60426 | 0.90716 |

FC, fold change; LEfSe, linear discriminant analysis effect size.

**Table S5. Bacterial genus abundance for KO vs WT.**

| **Genus** | **log2FC** | **LEfSe** | **p-value** |
| --- | --- | --- | --- |
| *Muribaculum* | 0.59113 | 0.16799 | 0.000434 |
| *Parasutterella* | 0.83072 | 0.25001 | 0.000892 |
| *Bacteroides* | 0.90825 | 0.31863 | 0.004365 |
| *Desulfovibrio* | 0.52296 | 0.24667 | 0.034 |
| *Odoribacter* | -0.60534 | 0.28856 | 0.035924 |
| *Turicibacter* | 2.3291 | 1.3631 | 0.087498 |
| *Prevotellaceae_UCG_001* | -0.45632 | 0.30679 | 0.13691 |
| *Helicobacter* | -0.64514 | 0.4853 | 0.18373 |
| *Ileibacterium* | 2.0864 | 1.8245 | 0.25282 |
| *Rikenella* | -0.52634 | 0.4802 | 0.27304 |
| *Alistipes* | 0.27247 | 0.28205 | 0.33402 |
| *Streptococcus* | 0.34098 | 0.50571 | 0.50015 |
| *Butyricicoccus* | -0.319 | 0.53422 | 0.55042 |
| *Colidextribacter* | 0.0762 | 0.12966 | 0.55674 |
| *Mucispirillum* | -0.15845 | 0.33957 | 0.64078 |
| *Lachnospiraceae_UCG_001* | -0.60117 | 1.3372 | 0.65302 |
| *Oscillibacter* | -0.13072 | 0.39974 | 0.74366 |
| *Rikenellaceae_RC9_gut_group* | 0.16419 | 0.60227 | 0.78515 |
| *Eubacterium_oxidoreducens_group* | 0.20478 | 1.0481 | 0.84509 |
| *Lachnospiraceae_UCG_006* | -0.17619 | 0.96157 | 0.85462 |
| *Blautia* | 0.12781 | 0.82583 | 0.877 |
| *Lachnospiraceae_NK4A136_group* | 0.020539 | 0.22245 | 0.92644 |
| *Parabacteroides* | 0.020913 | 0.23773 | 0.9299 |

FC, fold change; LEfSe, linear discriminant analysis effect size.

**Table S6**. **Pathways functionally associated with the microbiome analysis.**

tx-j vs C3H

| Pathway | Total | Expected | Hits | p-value |
| --- | --- | --- | --- | --- |
| Biosynthesis of amino acids | 222 | 12.7 | 44 | 3.81E-15 |
| Pyruvate metabolism | 74 | 4.23 | 18 | 6.17E-08 |
| Carbon fixation pathways in prokaryotes | 60 | 3.43 | 16 | 8.88E-08 |
| Nitrotoluene degradation | 7 | 0.4 | 6 | 2.07E-07 |
| Citrate cycle (TCA cycle) | 53 | 3.03 | 14 | 6.93E-07 |
| Carbon metabolism | 249 | 14.2 | 28 | 0.000216 |
| Butanoate metabolism | 61 | 3.48 | 10 | 0.00193 |
| Carbapenem biosynthesis | 2 | 0.114 | 2 | 0.00324 |
| Propanoate metabolism | 55 | 3.14 | 9 | 0.0033 |
| Valine, leucine and isoleucine biosynthesis | 15 | 0.857 | 4 | 0.00848 |
| Histidine metabolism | 35 | 2 | 6 | 0.0127 |
| Glycolysis / Gluconeogenesis | 80 | 4.57 | 10 | 0.0139 |
| 2-Oxocarboxylic acid metabolism | 57 | 3.26 | 8 | 0.0141 |
| Sulfur metabolism | 38 | 2.17 | 6 | 0.0188 |
| C5-Branched dibasic acid metabolism | 11 | 0.628 | 3 | 0.0214 |
| Arginine and proline metabolism | 115 | 6.57 | 12 | 0.0281 |
| Methane metabolism | 105 | 6 | 11 | 0.0341 |
| Terpenoid backbone biosynthesis | 23 | 1.31 | 4 | 0.0386 |
| Pyrimidine metabolism | 149 | 8.51 | 14 | 0.041 |

KO vs WT

| Pathway | Total | Expected | Hits | p-value |
| --- | --- | --- | --- | --- |
| Glycosaminoglycan degradation | 14 | 0.0567 | 3 | 1.64E-05 |
| Flavone and flavonol biosynthesis | 3 | 0.0121 | 1 | 0.0121 |
| Alanine, aspartate and glutamate metabolism | 62 | 0.251 | 2 | 0.0243 |
| Other glycan degradation | 7 | 0.0283 | 1 | 0.028 |
| Vitamin B6 metabolism | 12 | 0.0486 | 1 | 0.0476 |
| Biosynthesis of amino acids | 222 | 0.899 | 3 | 0.0524 |
| Valine, leucine and isoleucine biosynthesis | 15 | 0.0607 | 1 | 0.0592 |
| Nitrogen metabolism | 17 | 0.0688 | 1 | 0.0669 |
| Naphthalene degradation | 17 | 0.0688 | 1 | 0.0669 |
| Drug metabolism - cytochrome P450 | 19 | 0.0769 | 1 | 0.0745 |

**Table S7. Gross phenotypes of mouse models at 9 weeks of age.**

| **Genotype** | **n** | **Body weight (g)** | **Liver / Body weight** |
| --- | --- | --- | --- |
| WT chow | 3 | 21.4 ± 1.5 | 0.056 ± 0.002^a^ |
| KO chow | 3 | 20.1 ± 2.5 | 0.054 ± 0.004^a^ |
| WT HFD | 3 | 17.9 ± 0.4 | 0.045 ± 0.001^b^ |
| KO HFD | 4 | 19.6 ± 2.7 | 0.043 ± 0.005^b^ |
| iWT chow | 7 | 24.7 ± 4.3 | 0.049 ± 0.005^a^ |
| *Atp7b*^ΔIEC^ chow | 8 | 21.9 ± 2.4 | 0.056 ± 0.005^b^ |
| iWT HFD | 8 | 24.4 ± 4.3 | 0.045 ± 0.005^a^ |
| *Atp7b*^ΔIEC^ HFD | 8 | 21.9 ± 2.7 | 0.044 ± 0.003^a^ |

Values are means ± SD and statistical significance was determined by Student’s t-test with p<0.05 considered significant. Values with different letter symbols are significantly different from each other within the same mouse model (KO or *Atp7b*^ΔIEC^).

*Atp7b*^ΔIEC^, *Atp7b* intestine-specific knockout; chow, Purina LabDiet 5001; HFD, high-fat diet Research Diets D12492; iWT, Lox^+/+^/Cre^-^ control for *Atp7b*^ΔIEC^; KO, *Atp7b* null global knockout on C57Bl/6 background; WT, littermate controls (*Atp7b*^+/+^) for KO mice.
